# Rfx3 controls outer hair cell differentiation, maintenance, and hair bundle formation by regulating the expression of *Insm1, Ikzf2*, and *Triobp* genes

**DOI:** 10.1101/2024.09.24.614849

**Authors:** Penghui Zhang, Yafan Wang, Xiang Guo, Lu Ma, Xiangyao Zeng, Zhili Feng, Jinlei Liu, Mengzhen Yan, Yushan Gao, Jieran Dong, Junhong Li, Jie Ling, Hong Wu, Qianchen Jing, Yong Feng, Jun Li

## Abstract

The RFX family of transcription factors plays crucial roles in the regulation of ciliogenesis and organ development. Mutations of RFX transcription factors lead to various genetic diseases, including ciliopathies and hearing loss, although the underlying mechanisms remain unclear. This study comprehensively analyzed the expression patterns of RFX family members during different inner ear developmental stages. Rfx3 and Rfx7 were identified as the dominantly expressed members in cochlear hair cells, exhibiting alternative splicing variants and dynamic subcellular localization at different stages. Integration of single-cell transcriptomics, ChIP-seq, and ATAC-seq data indicates that Rfx3 functions as either a transcriptional activator or repressor, binding to numerous genes related to ciliogenesis, hair bundle structures, and planar cell polarity. Rfx3 regulates the spatiotemporal expression of hair bundle gene *Triobp* in hair cells by binding to and modulating its intronic enhancer. Additionally, Rfx3 binds to hair cell differentiation and fate determination genes *Tbx2*, *Insm1*, and *Ikzf2*. The results demonstrate that Rfx3 and Rfx7 are dominantly expressed in outer hair cells, with their subcellular localization shifting to cytoplasm at later developmental stages. This suggests a previously unknown function of Rfx3 and Rfx7 beyond transcriptional regulation, highlighting their complex roles in hair cell differentiation and maintenance.

## Introduction

The mammalian inner ear is a complex sensory organ that relies on specialized cells called hair cells to detect sound, movement, and orientation. Hair cells are distinguished by their “hair-like” cilia on the apical surfaces. The auditory hair cell has two types of cilia: kinocilia and stereocilia ^1^. Kinocilia are true cilia and exhibit the classic 9+2 organization of microtubules like a motile cilium. While kinocilia in mammals do not directly mediate auditory mechano-electrical transduction (MET), they are crucial for hair cell morphogenesis, planar cell polarity (PCP), and stereocilia organization. In mammals, the auditory kinocilium is a transient structure that degenerates post-mature ^2,3^. Stereocilia, in contrast, are not true microtubule-based cilia. Instead, stereocilia are composed of cross-linked filamentous actin (F-actin) ^1^. The mechanic ion channels are located at the top of stereocilia; their displacement opens these channels, facilitating MET transduction. Typically, a hair cell contains about 100 stereocilia on its apical surface, forming a hair bundle architecture. This structure is intricately organized and comprises core F-actin, actin regulatory and bundling proteins, motor proteins and mechano-transduction related proteins. This hair bundle acts as a mechanical switch, converting sound vibration into electrochemical neuronal signals ^1^.

Cilia are highly conserved and intricately organized organelles present across eukaryotes, from simple unicellular organisms to complex mammals. They are ubiquitously on various vertebrate cell types and play essential roles in the organogenesis, development, and neuronal sensation. Specialized cilia in sensory epithelial cells detect various signals, including fluid flow in the kidney, odorant molecules in olfactory sensory neurons, light in photoreceptor cells of the eyes, and sound and balance signals in inner ear hair cells ^2,3^. In vertebrates, ciliogenesis is tightly regulated at the transcriptional level, orchestrating cilia growth, assembly, morphogenesis, and maintenance. Mutations in ciliary components or associated factors are linked to a range of genetic disorders known as ciliopathies ^4^, affecting multiple organs or tissues, including the inner ear, eyes, olfactory system, central nervous system, respiratory system, kidney, liver, and muscle ^5^. A notable hearing-related ciliopathy is Usher syndrome, characterized by deafness, blindness, infertility, and movement anomalies. Furthermore, many human deafness genes are associated with hair bundle structure or function ^2,6,7^.

The regulatory factor X (RFX) family TFs are reported as the central regulators of ciliogenesis ^3,8^. RFX family TFs share a highly conserved winged-helix type DNA-binding domain that recognizes the DNA motif known as the X-box and they are commonly found in worms, flies, mice, and humans ^8^. Mammals possess eight RFX family members (*RFX1-8*), which are differentially expressed across various tissues or cell types. In addition to their DNA-binding domains (DBD), RFX TFs contain activation domain (AD), domain B, domain C, and dimerization domain (DIM)^9^. RFX TFs are known to function both as transcriptional activators and repressors ^10^, regulating ciliogenesis by binding to X-box motifs in the promoters or enhancers of target genes. RFX TFs can form homo- or heterodimers through their DIM domains ^10,11^. The RFX family members exhibit redundant functions, where the loss of one member’s function can be compensated by the expression of another ^12,13^.

Beyond ciliogenesis, RFX family TFs are involved in numerous functions, including organ development, immune response, and tumorigenesis. For example, Rfx2 regulates many cilia-related genes and testis-specific genes, with Rfx2-null male mice exhibiting sterility due to spermatogenesis arrest^14^. Apart from ciliogenesis, RFX3 is crucial for the development of the endocrine pancreas and the regulation of β-cell differentiation, function, and glucokinase expression ^15^. Rfx3 also plays a role in nodal cilium development and left-right asymmetry specification, with mutations causing high frequencies of left-right asymmetry abnormality and embryonic lethality^16^. Rfx4 mutant mice show aberrant ciliogenesis and Sonic Hedgehog signaling, resulting in spinal cord and brain developmental defects ^17^. Both RFX5 and RFX7 have essential roles in the immune system ^18,19^. Rfx5 was reported to regulate MHC class gene expression during immune responses ^18,20,21^. RFX7 is critical for cilia formation in neural tube development in *Xenopus laevis* ^22^. RFX7 was also reported to function as a ubiquitous regulator of cell growth and fate determination, involved in the p53 tumor suppressor network and tumorigenesis ^23^. Rfx7 coordinates a transcriptional network controlling cell metabolism, essential for natural killer lymphocytes maintenance and immunity ^19^. Mutations of RFX TFs are implicated in several human diseases, including Alstrom syndrome, Ciliary Dyskinesia, and hearing loss ^4,13^. The double knockout of Rfx1/Rfx3 results in progressive outer hair cell loss and hearing loss in adult mice^13^. Our previous publication reported that the master transcription factor Six1 can cooperate with Rfx1/3 to regulate critical gene expression during inner ear development ^24^. However, the detailed mechanisms regarding the target genes of RFX TFs, how they regulate ciliary gene expression, and whether they have redundant functions in auditory systems remain unclear.

Here, we investigated the temporal and spatial expression of RFX TFs in inner ears and found that Rfx3 and Rfx7 are the two main RFX family members dominantly expressed in cochlear hair cells. Both Rfx3 and Rfx7 displayed dynamic expression patterns and subcellular localizations at different developmental stages, suggesting their complex functions. By analyzing inner ear single-cell RNA-seq (scRNA-seq), Rfx3 chromatin immunoprecipitation sequencing (ChIP-seq), and publicly accessible ATAC-seq and ChIP-seq data ^24–26^, we systematically profiled the genome-wide binding features of Rfx3 and its target genes. Ciliogenesis-related genes were the most significantly enriched genes bound by Rfx3. Mouse transgenic reporter experiments confirmed that Rfx3 regulated the spatiotemporal expression of the hair bundle gene *Triobp* in hair cells by binding to an intronic cis-regulatory element (CRE). The transgenic analyses of CREs bound by Rfx3 in *Triobp*, *Myo1h*, and *Insm1* gene loci revealed that Rfx3 might act as either a transcription activator or repressor, depending on the DNA sequence context of these CREs and their binding transcription factors. Importantly, our ChIP-seq and transgenic reporter data, along with other public data, revealed that Rfx3 affects outer hair cell development, differentiation and maintenance by regulating *Ikzf2* and *Insm1* genes.

## Results

### Dynamic and differential expression of *RFX* family transcription factor genes in the inner ear

The Human (or mouse) RFX family TFs can be subdivided into three groups based on the features of their five functional domains. Group 1(RFX1-3) contains all typical domains; Group 2 (RFX4, 6, and 8) lacks the activation domain; Group 3 (RFX5 and RFX7) only has the DNA-binding domain (Figure 1A) ^9^. To understand the function of RFX genes in hearing, we first characterized the tissue or cell type expression of RFX family genes in the inner ear at different developmental stages. By utilizing our own mouse inner ear scRNA-seq data (submitted work, preprint version) ^27^ and public scRNA-seq data resources ^28^, we found that *Rfx3* and *Rfx7* were dominantly expressed in hair cells at different developmental stages of inner ears. At embryonic stage 16 (E16), the “hair cells” showed dominant expression of *Rfx3*, medium expression of *Rfx7*, and weak or no expression of other RFX family members (Figure 1C). The hair cells are not fully differentiated at the E16 stage, and the “hair cell progenitors” are in the pre-differentiation stage, having started to express hair cell markers. The *Rfx3* and *Rfx7* also showed certain expression in other cell types, such as sulcus cells (Figure 1C). At postnatal day 7 (P7), *Rfx3* and *Rfx7* were still the two dominant RFX genes expressed in hair cells in inner ears. (Figure 1D). Consistently, *Rfx3* and *Rfx7* remained dominantly expressed in hair cells of adult cochlea and were also strongly expressed in spiral ganglion neurons (SGNs) in adult mice. *Rfx3* and *Rfx7* also showed certain expression in other inner ear cell types (Figure 1E). Western blot analysis with an Rfx3 antibody from Sigma showed a decreased expression pattern from E13.5 to adult stages (Figure 1B, antibody 1). However, another Rfx3 antibody (Proteintech, antibody 2) displayed a relatively reversed expression pattern. These different expression patterns indicated the presence of different protein isoforms specifically recognized by these two antibodies. Similar to the Rfx3 Western blot results using antibody 1, the expression of Rfx7 decreased from E13.5 to adult stages (Figure 1B).

**Figure 1.**
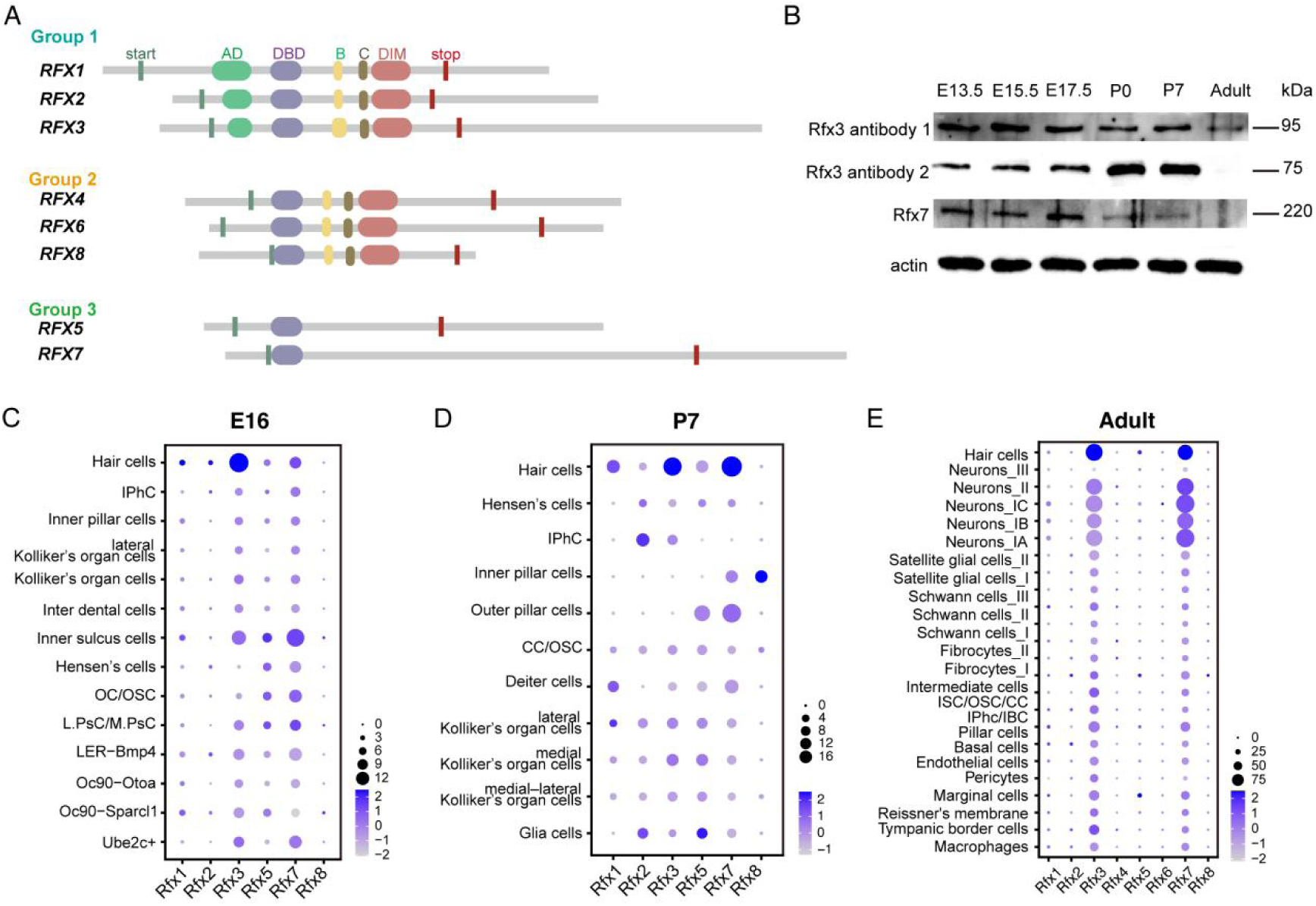
Expression of RFX family transcription factors in mouse inner ear single cell atlas. (A) Representative RFX family transcription factors are grouped according to functional domains. The first group consists of RFX1, RFX2, and RFX3, which have all the domains, including AD (activation domain), DBD (DNA binding domain), B (domain B), C (domain C), and DIM (dimerization domain). The second group consists of RFX4, RFX6, and RFX8, harboring the domains of DBD, B, C and DIM domains. The third group consists of RFX5 and RFX7, which only have the DBD. The green and red squares mark the start and stop codon positions, respectively. (B) Western blot with indicated antibodies in the whole cochlea at different developmental stages. Different developmental stages include E13.5, E15.5, E17.5, P0, P7, and adult (2M). (C,D,E) Single cell-level expression patterns of the RFX family TFs in three developmental stages in the mouse cochlea. IPhC: inner phalangeal cells/border cells, OC/OSC: Claudius cells/outer sulcus cells, L.PsC lateral prosensory cells, M.PsC medial prosensory cells, Oc90: Oc90+ cells, IPhc/IBC, inner phalangeal cell /inner border cell; ISC/OSC/CC: Claudius cells and inner and outer sulcus cells.

Immunostaining experiments were performed to further analyze the expression pattern of Rfx3 and Rfx7 as revealed by scRNA-seq data (Figure 2A-F). Rfx3 was dominantly expressed in all the hair cells at the E17.5 stage, both in the nucleus and cytoplasm (Figure 2A). However, the immunostaining signal of Rfx3 was dominantly observed only in outer hair cells (OHCs) rather than inner hair cells (IHCs) at the P7 stage. Additionally, their subcellular localization shifted to cytoplasm only (Figure 2B). Strikingly, immunofluorescent signals of Rfx3 became exclusively or dominantly expressed in IHCs in adults (Figure 2C). Besides, Rfx3 was strongly expressed in SGNs at all these stages (E17.5, P7, and adult) and in stria vascularis at the P7 and adult stages (Figure 2B and S1A). At the E17.5 stage, Rfx7 was ubiquitously expressed in many cell types throughout the cochlear duct and exclusively detected in the nucleus (Figure 2D). Relatively stronger expression of Rfx7 was observed in IHCs but not in OHCs. Similar to Rfx3, Rfx7 immunostaining signals were absent in IHCs and only detected in OHCs at the P7 stage (Figure 2E). Notably, Rfx7 proteins were distributed in both the nucleus and cytoplasm, distinct from their nuclear localization at the E17.5 stage and from the cytoplasmic localization of Rfx3 at the same stage (Figure 2A, B). Rfx7 was also strongly expressed in the stria vascularis and SGNs (Figure 2F,S1B,C) similar to Rfx3 and consistent with scRNA-seq data (Figure 1E). These immunostaining data revealed dynamic and varied expression patterns of Rfx3 and Rfx7 in the inner ear.

**Figure 2.**
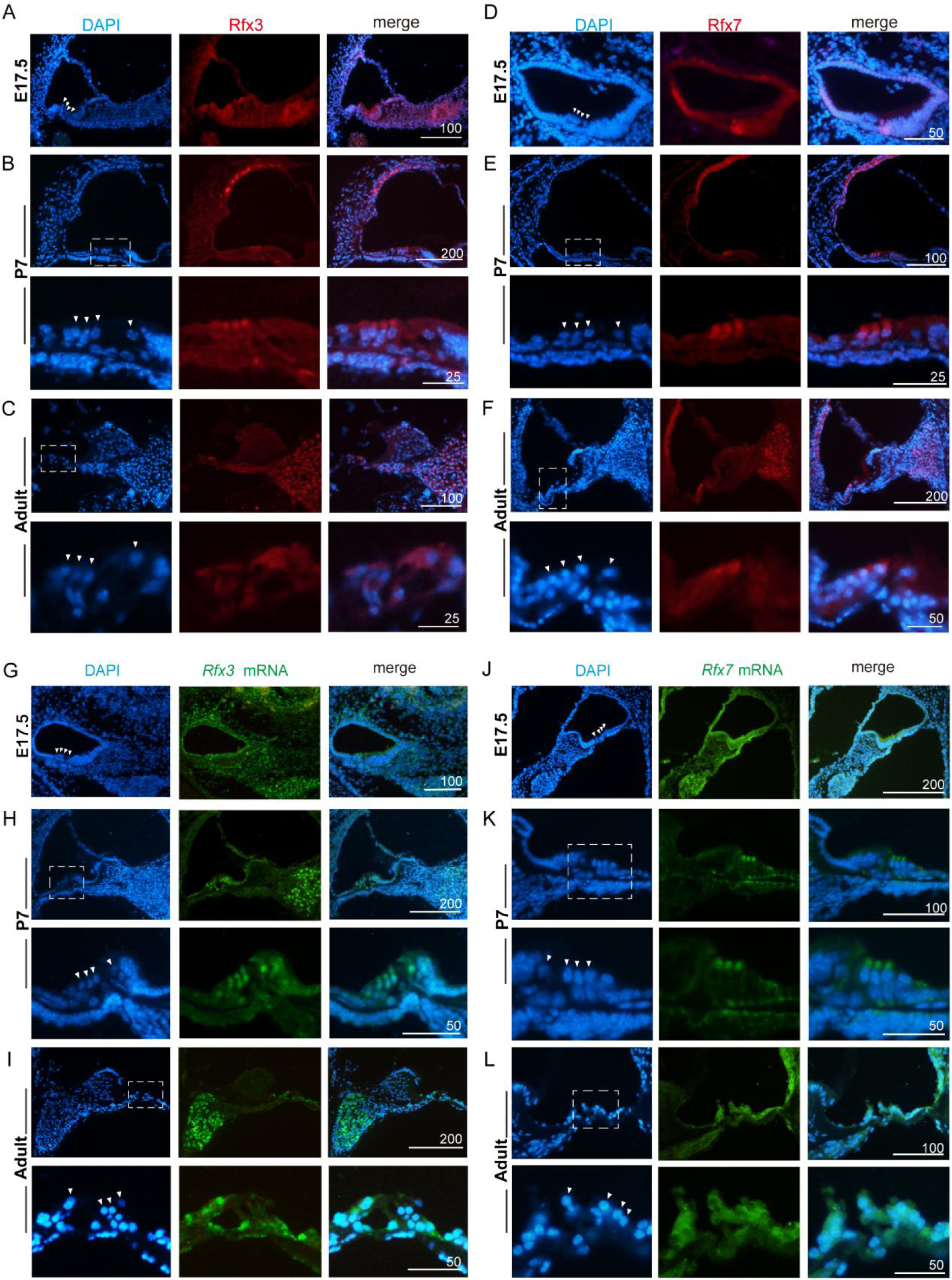
Spatiotemporal and dynamic expression patterns of Rfx3 and Rfx7 in the mouse inner ear. (A-C) Immunofluorescence detection of Rfx3 protein in organ of Corti at different developmental stages. All protein immunostaining signals are in red, nuclei were counterstained with DAPI (blue). Hereafter is the same. (A) Rfx3 expression on E17.5 cochlea section. (B) Rfx3 expression on P7 cochlea section. The white dashed boxed region was magnified and displayed below. Hereafter is the same. (C) Rfx3 expression on adult cochlea section. (D-F) Immunofluorescence detection of Rfx7 proteins in organ of Corti at different stages. (D) Rfx7 expression on E17.5 cochlea section. (E) Rfx7 expression on P7 cochlea section. (F) Rfx7 expression on adult cochlea section. (G-I) RNA *in situ* detection of *Rfx3* gene in organ of Corti at different developmental stages. All the RNA expression signals were labeled in green. The cochlea sections from E17.5 (G), P7 (H), and adult (I) wild-type stages revealed a dynamic expression pattern of Rfx3 gene during the development. (J-L) RNA i*n situ* detection of *Rfx7* gene on the cochlea sections. *Rfx7* showed dynamic expression patterns from E17.5 (J), P7 (K), and adult stage (L). The white triangles indicate the position of hair cells. Representative images from n>3 experiments. Scale bars are indicated in the figure (µm).

Further, RNA *in situ* detection of Rfx3 and Rfx7 genes was carried out on inner ear sections at different stages to confirm their complicated expression patterns in inner ears (Figure 2G-L). Consistent with the protein signals, *Rfx3* mRNAs were dominantly detected throughout the cochlear duct and SGNs at the E17.5 stage (Figure 2G). Remarkably, *Rfx3* mRNAs were ubiquitously found in all hair cells and were relatively more strongly expressed in IHCs at P7 (Figure 2H), differing from the protein immunostaining signals that labeled only OHCs (Figure 2B). Abundant *Rfx3* mRNAs were detected in SGNs and the stria vascularis, consistent with the protein detection pattern (Figure 2A,C). At the adult stage, strong *Rfx3* mRNAs were detected only in IHCs rather than OHCs, consistent with the protein immunostaining results (Figure 2I). Consistently, strong *Rfx3* mRNA signals were found in SGNs and stria vascularis. *Rfx7* mRNA detection displayed signal patterns similar to those of *Rfx3* mRNA (Figure 2J-L). Abundant *Rfx7* mRNAs were detected in the cochlear duct at the E17.5 stage (Figure 2J), but only in OHCs at P7 (Figure 2K) and adult stages (Figure 2L), consistent with the protein immunostaining results in hair cells (Figure 2D-F). Notably, *Rfx7* mRNAs were also detected in SGNs and the stria vascularis at all stages, consistent with Rfx7 antibody immunostaining signals in SGNs (Figure 2E,F,S1B,C).

Overall, the scRNA-seq data along with protein and mRNA level detection results consistently confirmed the dynamic expression pattern of Rfx3 and Rfx7 in major inner ear cell types. Some inconsistencies between protein immunostaining and RNA *in situ* detection results suggest the existence of splicing isoforms. For example, Rfx3 proteins were not dominantly detected in IHCs at the P7 stage (Figure 2B), but *Rfx3* mRNAs were found in all hair cells, with dominant expression in IHCs (Figure 2H). According to the company’s information, Rfx3 antibody 1 was raised against a partial peptide sequence (270-405 aa), while the Rfx7 antibody was raised against a very short peptide of the extreme C-terminal region of human RFX7 protein (between 1313 -1363 aa). Therefore, these inconsistencies may be due to the presence of different splicing isoforms of Rfx3 and Rfx7 that are specifically recognized by certain antibodies.

To understand the differences between the isoforms, the Rfx3 and RFX7 (human) protein isoforms were downloaded and aligned. The Rfx3 isoforms displayed variations at their N-terminal other than their C-terminal (Figure S2A). The alterative N-terminal variants may result in the presence or absence of nucleus localization signals, determining their subcellular localization. Mismatched sequences were observed in both the N- and C-terminal regions of aligned RFX7 isoforms (Figure S2B). Bulk RNA-seq data from E17.5, P8, and adult stage inner ears revealed the existence of alterative RNA splicing isoforms at different developmental stages (Figure S2C,D). Exon 2 of the *Rfx3* gene was not expressed at all stages. It appeared that several intronic peaks were expressed only at the P8 and adult stages. Exons 16 and 17 of *Rfx3* were weakly or not expressed at all stages (Figure S2C). Regarding the *Rfx7* gene, a peak was specifically observed in a region in the first intron close to the promoter at P8 and adult stage, but was absent at the E17.5 stage (Figure S2D). This observation is consistent with the alignment result of all the RFX7 isoforms available in NCBI, which showed that the 5’ and 3’ ends displayed alterative splicing variants (Figure S2A,B). Additionally, exon 3 of *Rfx7* showed a weaker peak at E17.5 than at P8 and adult data, while exon 7 of *Rfx7* showed a much stronger peak at E17.5 than at P8 and in adult stage (Figure S2D). Different isoforms of Rfx3 and Rfx7 may explain their subcellular localization transitions at different developmental stages and the inconsistency between immunostaining and RNA *in situ* results.

Collectively, the expression patterns of Rfx3 and Rfx7 were generally similar. They were dominantly expressed in hair cells, SGNs, and stria vascularis at all examined developmental stages (Figure 2,S1). Their protein subcellular localizations shifted to the cytoplasm at later stages, with their expressions becoming restricted to OHCs at certain stages (P7) (Figure 2B,E). These results suggest their similar or potentially redundant functions. The dynamic subcellular localization changes of Rfx3 or Rfx7 from the nucleus to the cytoplasm may also be caused by alterative splicing, resulting in the presence or absence of their nuclear localization signals.

### Dynamic and differential binding of Rfx3 in the auditory sensory epithelium

Given that *Rfx3* and *Rfx7* are the most abundantly expressed RFX family genes in the hair cells and SGNs of developing and adult inner ears, we aimed to further characterize their regulatory mechanism using ChIP-seq assays. We succeeded in performing ChIP-seq experiment with a commercial Rfx3 antibody and, therefore, focused mainly on the Rfx3 ChIP-seq data in this study. Only the E17.5 stage cochlear epithelia were used for Rfx3 ChIP-seq, as Rfx3 becomes cytoplasmic at later developmental and adult stages (Figure 2B,C).

Two replicates of Rfx3 ChIP-seq data were obtained, and the one with high quality and more peaks was chosen for further analysis. A total of 10,604 peaks were identified in the E17.5 stage Rfx3 ChIP-seq data, averagely distributed within 500 kb regions from the transcription start site (TSS) (Figure 3A). Rfx3 was found to mainly bind to the proximal promoter (∼39.17%), intronic (27.85%), and intergenic regions (27.91%) (Figure 3B). The binding of Rfx3 to DNA displayed a narrow sharp peak around the peak center within ±200 bp regions, a typical binding feature for a TF as expected (Figure 3C). The top enriched gene ontology (GO) terms of Rfx3 bound genes were related to pattern specification process, embryonic organ morphogenesis, cell fate specification, cytoskeleton organization, and cilium organization (Figure 3D). Interestingly, although fewer peaks were obtained from the replicate 2 data, the top gene-enriched GO terms were mainly related to cilium, including cilium organization, cilium assembly, and regulation of microtubule-based movement (Figure S3A). The high enrichment of cilium-related genes was consistent with the well-known function of RFX family TFs in ciliogenesis ^8^, confirming the high quality of our ChIP-seq data. All the Rfx3 bound peak sequences were further analyzed for motifs using HOMER ^29^. The top six most significantly enriched motifs were the RFX, Forkhead, MYB, NFAT, JUND, and SIX motifs (Figure 3E). The extraordinary high P value of RFX motif enrichment (P value = 1e-7708) further confirmed the quality of the ChIP-seq data. The SIX motif enrichment was consistent with our previous report that Rfx1/3 can interact with Six1 in co-immunoprecipitation experiments with mouse inner ear tissue^24^.

**Figure 3.**
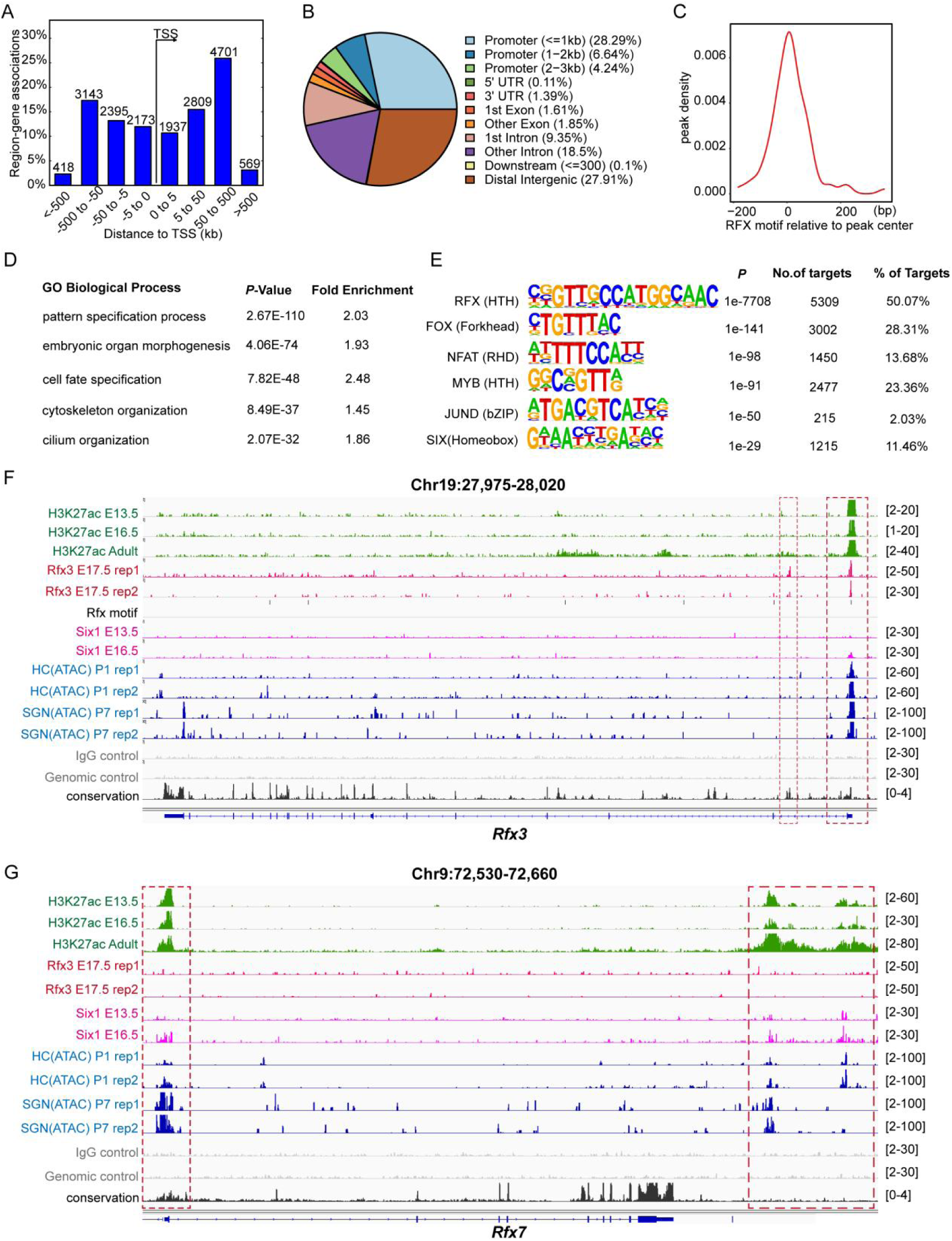
ChIP-seq analysis of Rfx3 in the mouse cochlea at E17.5 stage and *Rfx3* locus viewing of its regulatory features. (A) Distance of Rfx3 ChIP-seq peaks to TSS positions. (B) Genomic distribution of Rfx3-enriched regions. (C) Localization of the Rfx3-motif within the peak sequence revealed a typical binding feature of transcriptional factor. (D) The most enriched gene ontology (GO) terms of the Rfx3-bound genes. (E) The top 5 most enriched motifs in the Rfx3-bound CREs, were revealed by the Homer Known motif analysis. (F) *Rfx3* gene locus view revealed its regulatory features, including deposition of active marker H3K27ac, chromatin accessibility (ATAC-seq), and self-regulation by Rfx3. Two Rfx3-bound regions were indicated by red dashed box at the TSS and intron regions of *Rfx3* gene locus. (G) *Rfx7* gene locus view revealed its regulatory features, including H3K27ac deposition, chromatin accessibility, Rfx3 and Six1 binding.

Detailed locus views of *Rfx3* and *Rfx7* gene loci reveal their self-regulation and regulatory landscape. Strong enrichment of the activation marker H3K27ac at the promoter regions of Rfx3 and Rfx7 indicated that these genes were continuously expressed at E13.5, E16.5, and adult stages (Figure 3F,G). The RFX motif was clearly identified at the promoter of *Rfx3* gene. Rfx3 bound to its own promoter region and an intronic region to autoregulate its own gene expression (Figure 3F). However, Rfx3 did not occupy *Rfx7* locus. Similar to H3K27ac, chromatin accessibility as revealed by ATAC-seq is another critical marker of active gene expression. The ATAC-seq data of flow cytometry-sorted cochlear hair cells and SGNs ^25^ indicated their open access status and expression activity in these cells, consistent with scRNA-seq data and RNA *in situ* data (Figure 1, 2). Six1 was more enriched in the *Rfx7* locus rather than the *Rfx3* locus. The *Rfx7* gene locus has two potential enhancers that may coordinate its spatiotemporal expression in specific cell types, as revealed by their differential enrichment of ATAC-seq signals (Figure 3G).

### Rfx3-bound cis-regulatory elements can either be activated or silenced

Since Rfx3 has both transcriptional activating and repressive domains (Figure 1A), we further compared our Rfx3 ChIP-seq data with public SGN and hair cell ATAC-seq data and H3K27ac ChIP-seq data to reveal the regulatory features of CREs bound by Rfx3. Among the total 10,604 Rfx3 peaks, 1,543 peaks overlapped with H3K27ac-enriched regions, suggesting the activating regulation of Rfx3-bound CREs (Figure 4A, up). Clustered heatmaps confirmed different distribution patterns of H3K27ac signals near the Rfx3 peak center (Figure 4A, bottom). The ATAC-seq data of flow cytometry-sorted cochlear hair cells and SGNs ^25^ identified 50,070 and 39,778 peaks, respectively. Accordingly, 2,979 and 2,501 common peaks were identified between Rfx3 and ATAC-seq peaks, respectively (Figure 4B,C). The overlap of Rfx3 with both H3K27ac and ATAC-seq signals confirmed its activation role in gene expression regulation. Comparison and colocalization analysis of Rfx3, H3K27ac, and ATAC-seq peaks showed differential occupation features of these peaks. Seven clusters, corresponding to the Venn diagram, demonstrated differentiated features of Rfx3-bound CREs (Figure 4D). Cluster A Rfx3-bound CREs showed the absence of both ATAC-seq and H3K27ac signals (Figure 4D,S4B), indicating their repressive status in hair cells or SGNs. While cluster G CREs (Figure 4D), which showed the presence of all these signals, were in an active status.

**Figure 4.**
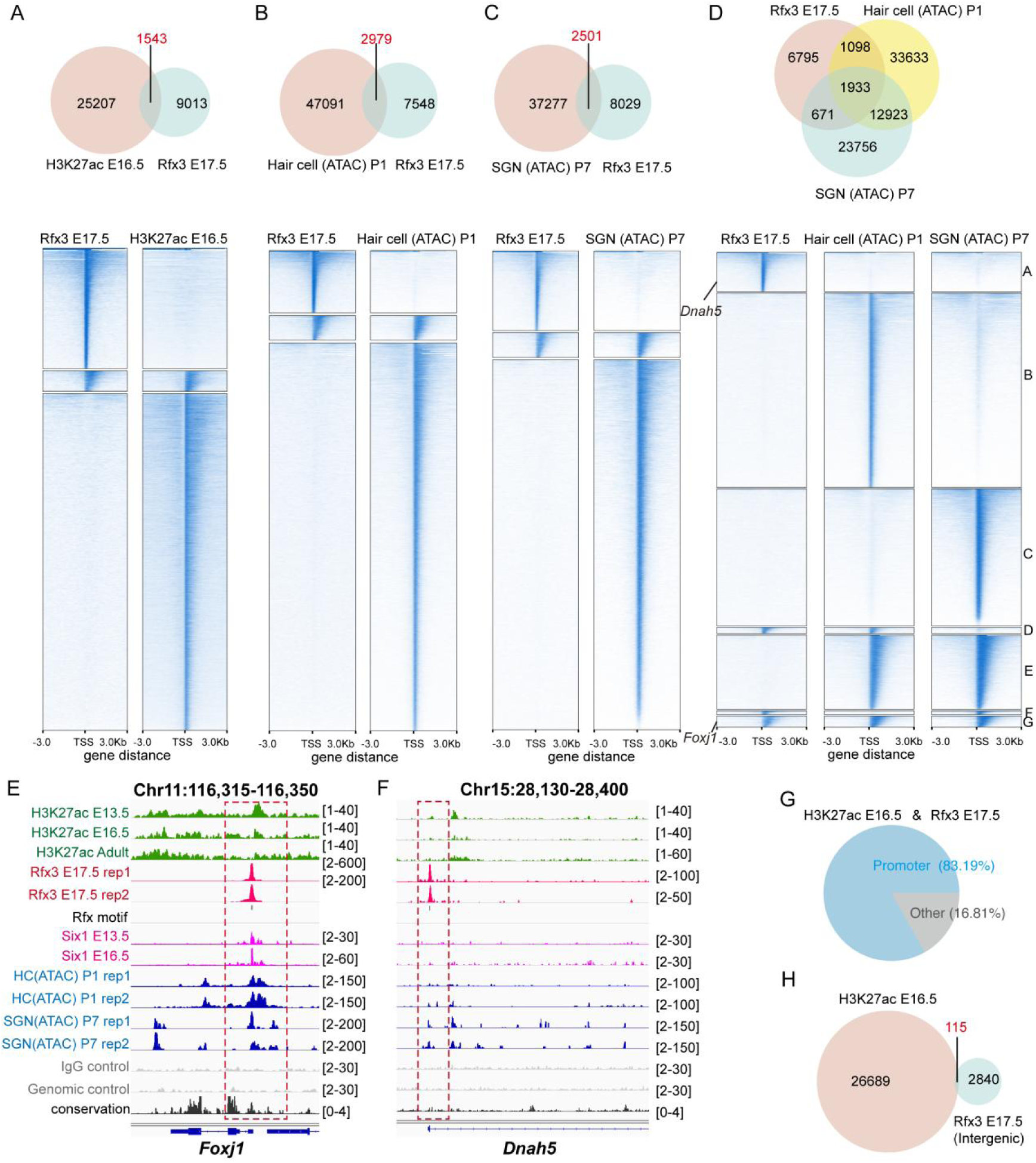
Rfx3 may act as either a transcriptional activator or repressor. (A) The Venn diagram of 1543 common peaks between the E17.5 Rfx3 bound cis-regulatory elements (CREs) at E17.5 and E16.5 H3K27ac enriched regions. (B) There are 2979 common peaks between Rfx3 and hair cell specific ATAC-seq peaks. (C) Venn diagram showed 2501 common peaks between Rfx3 and spiral ganglion neuron (SGN) specific ATAC-seq peaks. (D) Comparisons between E17.5 Rfx3 ChIP-seq, P1 hair cell ATAC-seq, and P7 SGN ATAC-seq peaks together. Clustered heatmaps of Rfx3, H3K27ac, Hair cell ATAC-seq and SGN ATAC-seq peaks respectively within a -3 kb/+ 3 kb window. (E) Strong enrichments of Rfx3, Six1, and ATAC-seq signals in the promoter of *Foxj1* genes. (F) Strong binding of Rfx3, however, no obvious deposits of H3K27ac, Six1 and Rfx3 in *Dnah5* gene locus. (G) The large majority of 1543 common peaks (A) between Rfx3 and H3k27ac locate in or around the promoter region. (H) The Venn diagram shows that most of 2955 intergenic Rfx3 bound CREs do not have H3K27ac signals.

For example, the *Foxj1* gene locus, a well-known downstream target of RFX TFs ^30^, showed strong binding of Rfx3 at its promoter region, along with H3K27ac and ATAC-seq signal enrichments (Figure 4E). In contrast, the *Dnah5* gene locus showed no enrichment of H3k27ac or ATAC-seq signals, despite strong enrichment of Rfx3 at its promoter region (Figure 4F). These results suggest that the binding of Rfx3 may have different regulatory outcomes regarding gene expression. Among the 1,543 common peaks between Rfx3 ChIP-seq and H3K27ac peaks, about 83.19% of peaks were in promoter or proximal promoter regions (Figure 4G). A few common peaks were located in intronic and intergenic regions (16.8% in total). For the 2,955 intergenic regions bound by Rfx3, only 115 peaks had ATAC-seq signal enrichment, while most lacked ATAC-seq peaks (Figure 4H), suggesting that the intergenic Rfx3 peaks mainly function as repressive regulatory elements. Therefore, the activating or repressive roles of Rfx3 may be context-dependent. When Rfx3 binds to promoter regions, it tends to act as a transcription activator. Conversely, intergenic binding of Rfx3 results in transcriptional repression.

### Rfx3 regulates a large number of genes associated with ciliogenesis in the inner ear

The analyses of Rfx3 peak-associated genes revealed that Rfx3 regulates a series of ciliogenesis related genes (Table S1), consistent with its well-known function in controlling ciliogenesis. We found that Rfx3 binds to the promoters of many intraflagellar transport (IFT) genes, including *Ift22*, *Ift27*, *Ift43*, *Ift52*, *Ift57*, *Ift74*, *Ift80*, *Ift81*, *Ift88*, *Ift122*, *Ift140*, and *Ift172* (except *Ift20* and *Ift46*) (Table S1). Rfx3 also binds to several centrosome or basal body related genes like *Nphp1*, *Bbs1*, and *Alms1*, and other cilia genes including *Kif3a*, *Dnah5*, *Pkd1*, and *Smo*. Reanalysis of scRNA-seq data revealed the expression of typical ciliogenesis-related gene in the inner ear cell atlas (Figure S5A). Most of these genes were not specifically expressed in hair cells. However, the type I and II neurons commonly expressed relatively higher levels of ciliogenes genes, such as *Kif3b*, *Ift88*, *Pkd1*, *Bbs2/4/5/7*, and *Alms1*. In the *Alms1* locus, strong Rfx3 signals, along with enriched H3K27ac signals, were consistently found at the promoter regions at different developmental stages (Figure S5B). The TF Six1 was also enriched at the promoter region. Differential chromatin open access features were shown in cochlear hair cells and SGNs, consistent with the gene expression feature of *Alms1* in the single-cell atlas (Figure S5B). At the *Ift172* gene locus, the enrichments of H3K27ac, Rfx3, and ATAC-seq signals were consistent with the expression features of *Ift172* in hair cells and SGNs (Figure S5C). Furthermore, Rfx3 may bind to the promoter of the *Ift81* gene (Figure S5D) to regulate its expression in hair cells (Figure S5E).

### Rfx3 regulates hair-bundle related gene *Triobp* and *Myo1h* in hair cells

Besides the top significant terms related to ciliogenesis, hair-bundle-related terms were also enriched in the GO analyses of Rfx3-bound genes (Figure S3A). A large number of hair-bundle-related genes were bound by Rfx3, as listed in Table S2. To investigate the regulatory functions of Rfx3 for hair bundle architecture, the expression profiles of hair-bundle-related genes in the single-cell atlas were analyzed to confirm their specific expression patterns in hair cells. Several hair-bundle-related genes, including *Myo7a*, *Cdh23*, *Triobp*, and *Ush1c*, displayed specific expression exclusively in hair cells (Figure 5A). In the *Myo7a* locus, Rfx3 bound to an intronic regulatory element in the upstream region of the gene (Figure S6A). This region only displayed open chromatin feature in hair cells rather than SGNs, as indicated by the presence of ATAC-seq enrichment exclusively in hair cells. The result suggests that this region is a cis-regulatory element (CRE) determining the exclusive expression pattern of *Myo7a* in hair cells but not in other cell types (Figure S6A). Rfx3 also bound to several regions of the *Cdh23* gene (Figure S6B), which is essential for formation of the tip link in the hair bundle ^6^. Multiple bindings of Rfx3 to the CREs of the *Cdh23* gene suggest a critical role for Rfx3 in hair bundle architecture and formation.

**Figure 5.**
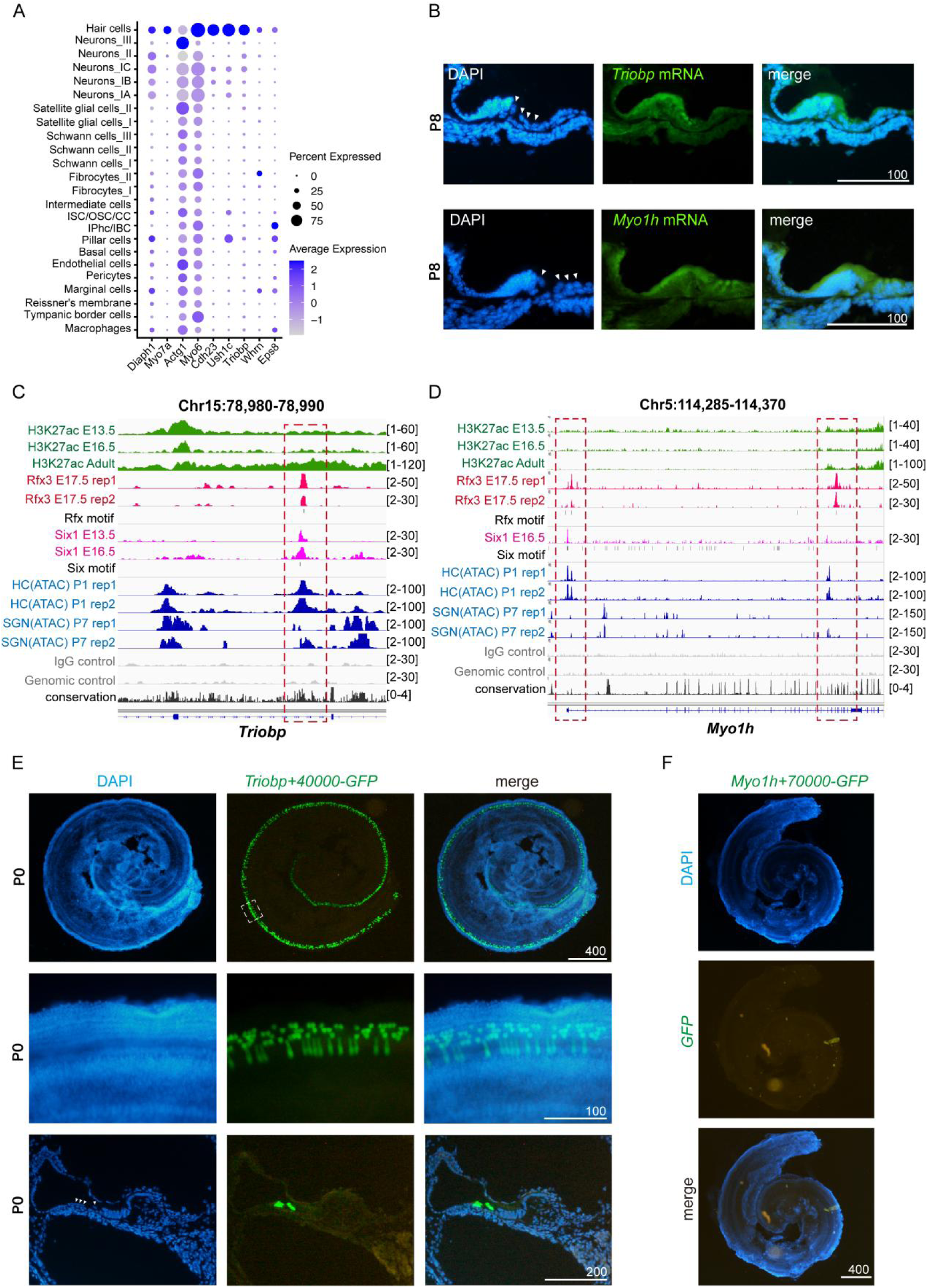
Rfx3 regulates the spatiotemporal expression of hair bundle related genes. (A) Single-cell level profiling of hair bundle related gene expresses in the adult cochlea. (B) RNA *in situ* detections of *Triobp* (upper panels) and *Myo1h* (lower panels) genes on cochlear sections in organ of Corti at P8. The *Triobp* and *Myo1h* probes were labelled in green, and nuclei were counterstained with DAPI in blue. The triangles indicate hair cells). (C) Genome browser visualization of Rfx3 peaks at the intron of *Triobp* gene. The RFX and SIX motifs were seen around the peak center. The red dashed box indicates Rfx3 bound regions. (D) Genome browser visualization of Rfx3 peaks at the TSS and intronic regions of *Myo1h* (E) Transient (G0) transgenic analysis of a 965-bp Rfx3-bound CRE (the red-dashed-boxed-region in Figure 5C) in *Triobp* gene at P0 stage. This +40 kb intronic CRE (*Triobp*+40000) drives specific expression of GFP reporter in hair cells (in 4/4 transgenic lines). Top panels, images of whole cochlea; middle panels, higher magnification of the partial areas in the top panel; bottom panels, images of cochlear sections showing GFP+ hair cells in the organ of Corti. (F) Transient (G0) transgenic analysis of an 847-bp Rfx3-bound CRE in the *Myo1h* intronic region (the red-dashed-boxed-region in Figure 5D, right) at P0 stage. The +70 kb intronic CRE (*Myo1h*+70000) did not show expression activity in P0 cochlea (5/5, n=5 transgenic lines). Scale bars are indicated in the figure (µm).

The actin-bundling protein Triobp is crucial for rootlet formation in hair cells and is essential for hearing ^31^. The RNA *in situ* detection of the *Triobp* gene confirmed its specific expression in hair cells, as revealed by scRNA-seq data (Figure 5A,B). *Triobp* showed stronger expression in OHCs. The H3K27ac and ATAC-seq peaks in the *Triobp* locus confirmed its active expression status in the inner ear (Figure 5C). The differential ATAC-seq peak features in the upstream gene region in hair cells and SGNs suggested the presence of alternative promoters in these two cell types, consistent with previous report indicating several splicing isoforms of the *Triobp* gene in the inner ear (exons 1 and 4 harbor two alternative promoter sites) ^31^. The isoform with a promoter located at exon 4 seemed to be the dominant *Triobp* variant in hair cells (Figure S5F, as indicated by red dashed box). The downstream Rfx3-bound region (red dashed box) showed colocalization of Rfx3 with Six1, H3K27ac and hair cell ATAC-seq peak (Figure 5C), suggesting this CRE region as a potential enhancer driving the specific expression of *Triobp* in hair cells. Subsequently, this 965-bp CRE region was cloned into the enhancer GFP reporter vector ^24^ to test its spatiotemporal specificity in gene expression regulation in transgenic mice. As expected, this CRE drove a specific expression pattern of GFP exclusively in hair cells in the transgenic mouse reporter experiment (4/4, n=4) (Figure 5E), confirming it as an enhancer activating the expression of the *Triobp* gene. Although GFP displayed a mosaic expression pattern in hair cells, this is normal for pronuclear injection due to DNA integration at relatively later-stage embryos. Overall, the spatiotemporal expression specificity of this enhancer is consistent with its endogenous specific expression pattern in hair cells, as revealed by scRNA-seq and RNA *in situ* data (Figure 5A,B).

For another hair-cell-specific expression gene, *Myo1h*, its function in the hair bundle and in hearing has not been studied and reported yet. Myo1h belongs to the class I myosins ^32^ and is predicted to be correlated with actin filament binding and microfilament motor activity. The ChIP-seq data revealed obvious enrichments of Rfx3 near the promoter and downstream regions (Figure 5D). The promoter or adjacent region showed enrichment of Rfx3 and ATAC-seq signals in hair cells only, suggesting that it was an activate regulatory region in hair cells but not in SGNs. The downstream intronic strong Rfx3 peak did not colocalize with ATAC-seq or H3K27ac signals in hair cells. Therefore, this Rfx3-bound CRE may be a repressive regulatory region (Figure 5D). To further confirm this, we cloned this 847-bp intronic Rfx3-bound CRE region into the reporter vector and analyzed its regulatory function in the hair cells of transgenic mice. All the positive transgenic embryos consistently showed no expression of specific GFP or very weak background signals (5/5, n=5) (Figure 5F), suggesting that this Rfx3-bound CRE was probably a repressor. The weak or absence of H3K27ac and ATAC-seq peak signals, along with the negative expression of the transgenic reporter, collectively revealed and validated the repressive function of Rfx3.

### Cooperation of Rfx3 with Forkhead-box transcription factors in the regulation of inner ear development

Transcription factors (TFs) normally function cooperatively. Our previous study reported that Six1 can interact with Rfx1 or Rfx3 to regulate the expression of several genes critical for inner ear development ^24^. There were 740 common peaks between Rfx3 and Six1 ChIP-seq data (Figure 6A). The SIX motif was also enriched in Rfx3 peak sequences (Figure 3E) and in the 740 Six1/Rfx3 co-bound CREs (Figure 6B), consistent with our previous findings that the RFX motif was identified in Six1-bound CREs ^24^. The mutual identification of RFX and SIX motifs from each other’s bound CREs confirmed their cooperative roles in gene regulations. Notably, the Forkhead transcription factor binding motif was one of the most significantly enriched motifs in the motif analysis of Rfx3-bound peak sequences (Figure 3E). Both Forkhead and SIX TFs were reported to physically interact with RFX TFs ^24,30^. These results suggest that their cooperative roles can be mediated either by their physical interaction at the protein level or by the presence of both motifs in a single CRE. The presence of these TF motifs and their sequence organization may facilitate or enhance the protein level interactions, similar to how the RFX motif dimer enhances the RFX-RFX protein self-interaction mediated by the DIM domain.

**Figure 6.**
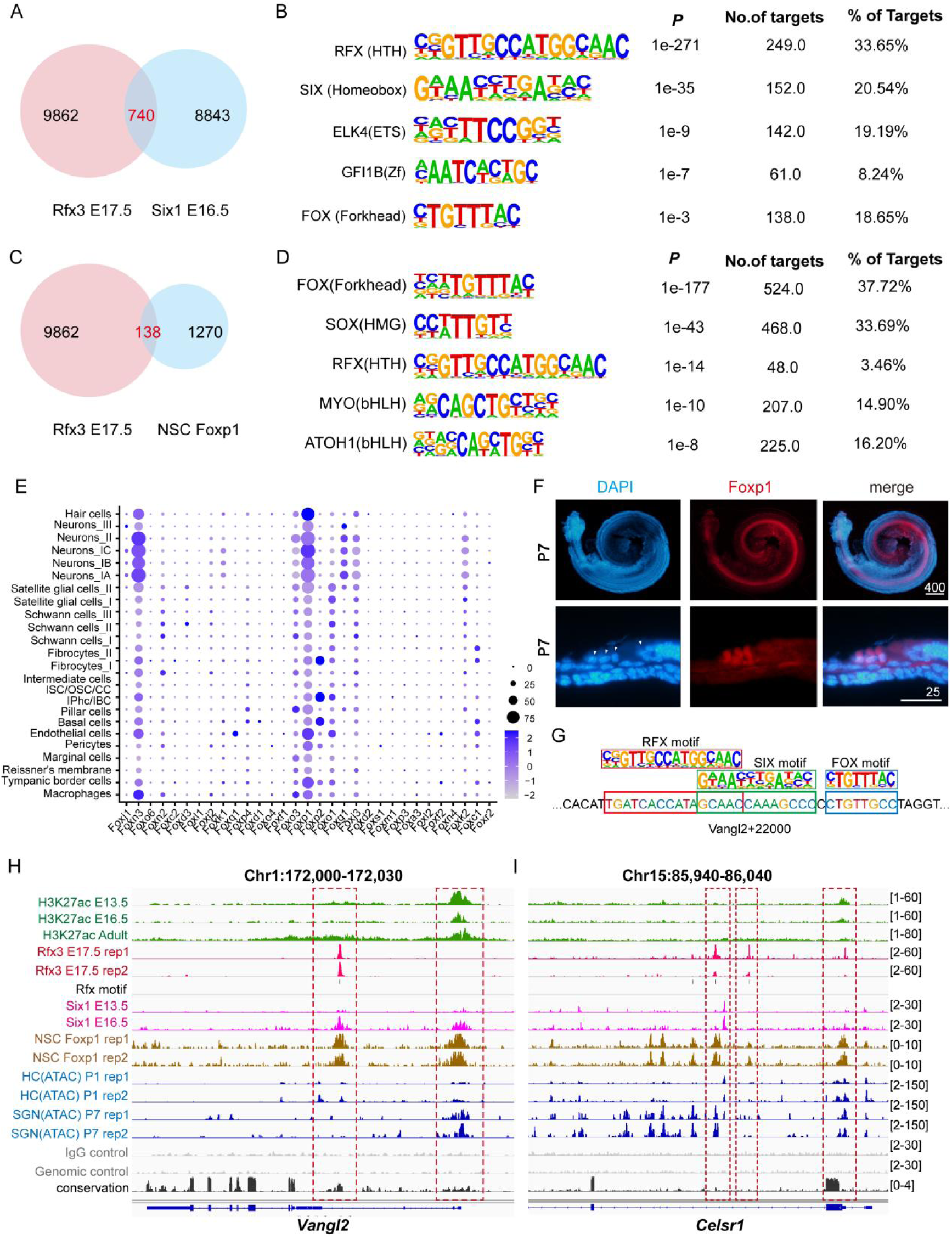
Rfx3 cooperates with Six1 and Foxp1 to regulate the planar cell polarity gene expression. (A) The Venn diagram shows that there are 740 common CREs bound by Rfx3 (E17.5) and Six1 (E16.5). (B) SIX motif was one of the top 5 most enriched motifs in the 740 co-bound CREs, as revealed by the Homer Known motif analysis. (C) The Venn diagram shows 138 common peaks between Rfx3 and Foxp1 binding CREs in neural stem cells (NSC). (D) Homer motif analysis of 138 co-bound peaks identifies RFX as one of the most enriched motifs in Foxp1-bound CREs. (E) Single-cell level profiling of the Forkhead family TFs gene expressions in the adult cochlea. (F) The whole amount and section of immunofluorescence showing the expression pattern of Foxp1 in organ of Corti (P7). Top panels, whole cochlea duct. Bottom panels, images of cochlear sections. (G) *Vangl2+22000* enhancer contains RFX, SIX and FOX-motifs. (H) Genome browser visualization of Rfx3 peaks at the TSS and intron of *Vangl2*, as indicated by the red dashed boxes The tested CRE region is highly conserved as revealed by the conservation score track (black, bottom). (I) Genome browser visualization of Rfx3 peaks at the TSS and intron of *Celsr1.* Three Rfx3-bound regions were indicated by the red dashed boxes. Scale bars are indicated in the figure (µm).

Forkhead motifs and other top-enriched TF motifs correspond to the top potential Rfx3-interacting TFs. To identify which Forkhead TFs are specifically expressed in hair cells or SGNs, the expression profiles of most Forkhead TFs in a single-cell atlas were characterized (Figure 6E). Foxn3 and Foxp1 were the two Forkhead TFs abundantly expressed in hair cells and SGNs, with similar expression patterns to Rfx3 and Rfx7. Different cell types seemed to specifically express certain Forkhead TFs. The Foxp1 antibody immunostaining of the whole amount cochlea epithelia and cryosection confirmed the expression pattern of Foxp1 revealed by single-cell data (Figure 6E,F). Foxp1 displayed strong expression in OHCs, similar to the expression pattern of Rfx3. Therefore, the public Foxp1 ChIP-seq data were analyzed to further validate the cooperativity between Rfx3 and Foxp1 ^26^. There were 138 common peaks between Rfx3 and Foxp1 ChIP-seq data (Figure 6C). Interestingly, the RFX motifs were highly enriched in Foxp1-bound CREs, as revealed by the motif analysis (Figure 6D). The mutual identification of both RFX and Forkhead motifs from each other’s CREs confirmed their cooperative roles, similar to the cooperation between RFX and SIX TFs. Consistently, Rfx3 peaks were exactly colocalized with Foxp1 peaks in many gene loci. For example, at the PCP gene *Vangl2* locus, colocalization of H3K27ac, ATAC-seq, Rfx3, Six1, and Foxp1 peak signals were observed at an intronic enhancer region (Figure 6H). Consistently, typical RFX, SIX, and Forkhead motifs were identified in this CRE sequence (Figure 6G,S6C). This enhancer was reported to drive specific expression of *Vangl2* in inner hair cells and surrounding supporting cells ^24^. These results suggest that all these TFs physically interact and function cooperatively to control the spatiotemporal expression pattern of the *Vangl2* gene by regulating this intronic enhancer activity. Similarly, Rfx3, Six1, and Foxp1 cooperate to regulate another PCP gene, *Celsr1* (Figure 6I). These data suggest that Rfx3 is essential for PCP signaling during the inner ear development, consistent with previous reports that Rfx3 is involved in PCP signaling and left-right axis development ^3,16^.

### Rfx3 is essential for hair cell differentiation, maturation, and maintenance

In addition to its role in PCP genes, Rfx3 was found to bind to several critical genes involved in hair cell differentiation, fate determination, and maintenance, including *Tbx2*, *Insm1*, *Ikzf2*, and *Barhl1* ^13,33^. Previous study revealed that Rfx1/3 double cKO mice underwent significant progressive OHC loss starting from P12 and lost all their OHCs throughout the length of cochlear duct by P90, although the OHCs differentiation, hair bundle structure, and PCP were not affected in the early stage ^13^.

To investigate the regulatory roles of RFX family TFs in OHC/IHC cell differentiation and maintenance, we first profile the specific expression patterns of each RFX TF family member and the genes *Tbx2*, *Insm1*, *Ikzf2* and *Barhl1* in IHCs and OHCs (Figure 7A). At E16, Rfx3 was mainly expressed in “IHCs” precursor cells, while Rfx7 was mainly expressed in “OHC” precursor cells. *Insm1* gene was dominantly expressed in the “OHC” cells, and *Tbx2* was mainly expressed in “IHC” cells. At the P7 stage, Rfx3 and Rfx7 proteins were mainly expressed in OHCs. The expression of *Insm1* ceased, and *Ikzf2* became strongly expressed in OHCs, consistent with the expression pattern of *Insm1* and *Ikzf2* in previous publication ^34^. *Tbx2* remained exclusively expressed in IHCs as expected at all stages. Strikingly, the expression pattern of Rfx3 and Rfx7 became reversed at the adult stage (Figure 7A). This expression pattern of Rfx3/7 in IHC/OHC at the P7 and adult stages in single-cell data showed certain inconsistences with RNA *in situ* and immunostaining results using the Rfx3/7 antibodies (Figure 2B,E).

**Figure 7.**
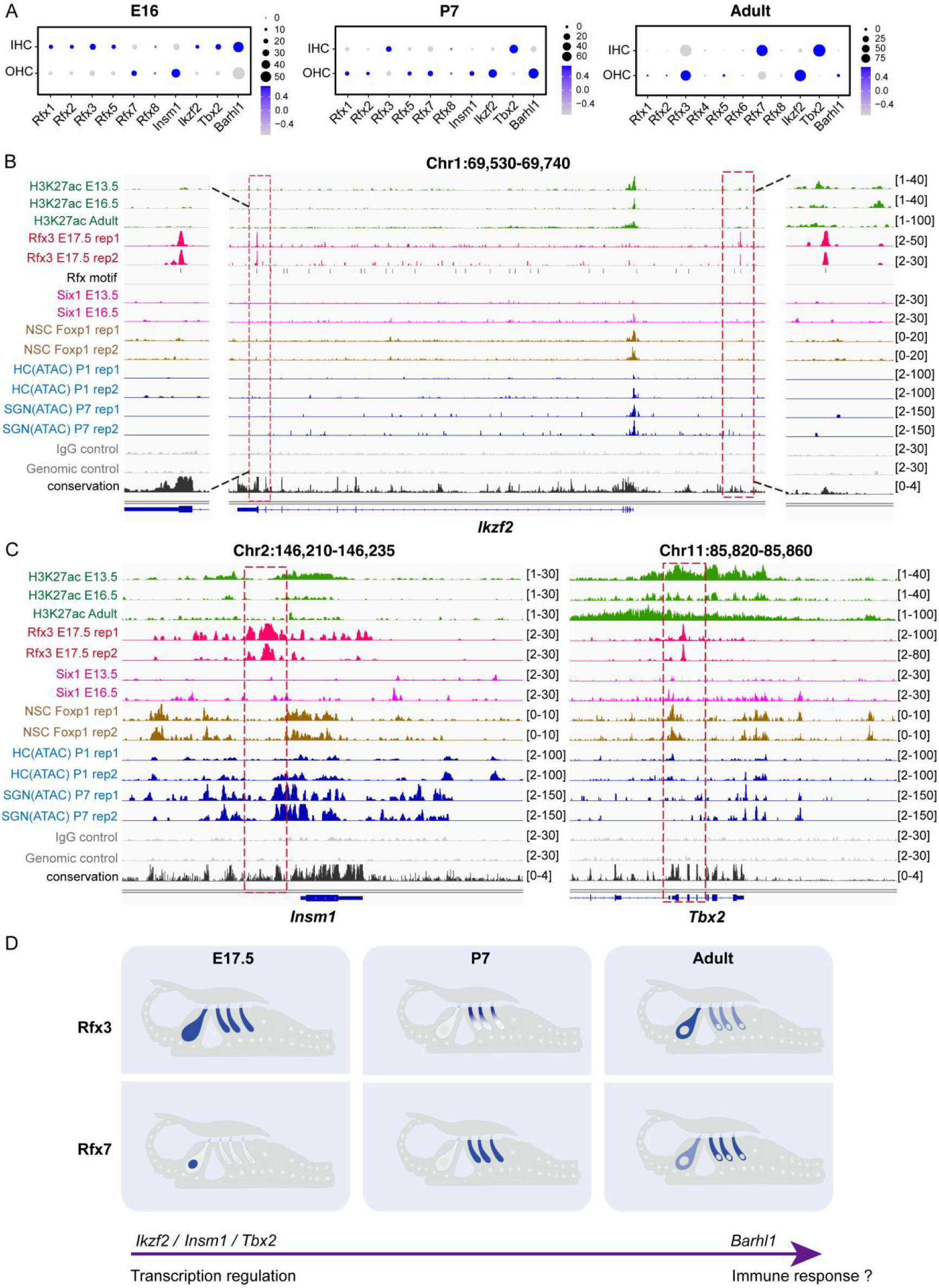
Rfx3 regulates *Tbx2*, *Insm1*, and *Ikzf2* gene expression and plays critical roles in hair cell differentiation and maintenance. (A) Expression of RFX TFs, and hair cell differentiation and maintenance genes at E16, P7 and adult stages in inner ears. (B) Genome browser visualization of Rfx3 peaks at the *Ikzf2* locus. Two Rfx3-bound regions were seen in the upstream intergenic and 3’ TTS regions. Left panels, higher magnification of the areas indicated by dashed lines at TSS; middle panels, whole *Ikzf2* locus view; right panels, higher magnification of the upstream intergenic region. The Rfx3 bound CREs were indicated by the red dashed box3s. (C) Genome browser visualization of Rfx3 peaks at the CREs of *Insm1* and intron of *Tbx2*. (D) The schematic figure describing the dynamic expression pattern of Rfx3 and Rfx7 during inner ear development (left, E17.5; middle, P7; right, adult). The arrows indicate a shift from the transcriptional regulatory role towards a possible immune response function.

We further analyzed Rfx3 ChIP-seq data to investigate the regulatory mechanisms of Rfx3 on its target genes *Ikzf2*, *Insm1*, *Tbx2*, and *Barhl1*. At the *Ikzf2* gene locus, the Rfx3 bound to an upstream intergenic region and the last downstream exon (Figure 7B). The *Ikzf2* gene was mainly expressed in OHCs and supporting cells (Figure 7A,S7A). Neither H3K27ac, ATAC-seq, nor Six1 signals were present in these two Rfx3-bound regions, suggesting these CREs were silenced and that Rfx3 likely acts as a silencer for *Ikzf2*. At the *Insm1* gene locus, Rfx3 bound to a region adjacent to a high ATAC-seq signal-enriched region (Figure 7C), suggesting Rfx3 may silence this active regulatory element. This 711-bp Rfx3-bound CRE was examined by transgenic reporter assay, and no expression activity was detected in any positive transgenic embryos (8/8, n=8) (Figure S7C), confirming the repressive role of Rfx3 in the *Insm1* locus. Similarly, the *Tbx2* gene locus showed strong enrichment of Rfx3 in an intronic region. However, this region lacked ATAC-seq signals in hair cells and SGNs, suggesting the repressive regulatory role of Rfx3 on *Tbx2* gene. Typical RFX, SIX, SOX and GATA motifs were identified within the Rfx3-bound CREs in *Ikzf2* and *Insm1* genes, suggesting their cooperative roles in silencing of these CREs (Figure S7D,E). Additionally, Rfx3 showed strong enrichment in the promoter region of the *Barhl1* genes (Figure S7B), suggesting Rfx3 regulates expression of *Barhl1* gene and is involved in hair cell survival. These results reveal that Rfx3 is highly correlated with hair cell differentiation and OHC maintenance by regulating *Tbx2*, *Insm1*, *Ikzf2*, and *Barhl1*.

## Discussion

### Compound functions of Rfx3 and Rfx7 in the inner ear

This study revealed a dynamic localization feature of Rfx3 and Rfx7 in inner hair cells (IHCs) or outer hair cells (OHCs) during different developmental stages. The transition of their subcellular localization from the nucleus to the cytoplasm indicates a functional shift from early developmental stages to adulthood. Besides the well-known transcriptional regulatory roles of RFX family members in ciliogenesis, they are reported to play crucial roles in organ development, PCP, left-right asymmetry specification, cell differentiation, and cell fate determination. Importantly, RFX TFs are essential for cell homeostasis and immunity ^35^. It has been reported that Rfx1/3 cKO mice progressively lose their OHCs starting from P12 and lose all OHCs in adulthood ^13^. After P7 or in adult stages, our data revealed that Rfx3 and Rfx7 translocated to cytoplasm (Figure 2). This cellular colocalization aligns with the deficiency phenotypes observed in mutant mice. These data suggest that Rfx3 no longer exerts transcriptional roles in the nucleus but instead translocates to the cytoplasm, potentially switching to a role in immune response, which is critical for hair cell homeostasis and maintenance. The faster and more severe loss of OHCs compared to IHCs may be due to the progressive loss of Rfx3 and Rfx7 expressions in OHCs in adulthood ^13^. These findings suggest that Rfx3/7 may play at least dual roles in the inner ears. A schematic model figure was proposed to illustrate the complex functions of Rfx3 and Rfx7 in hair cells (Figure 7D).

At earlier developmental stages, Rfx3/7 regulate the transcription of numerous genes related to ciliogenes or hair bundles in the nucleus. Specifically, Rfx3 may regulate TFs critical for hair cell development, fate determination, and differentiation, including *Tbx2*, *Insm1*, and *Ikzf2* genes. The regulation of these genes is consistent with the dynamic localization feature of Rfx3/7 in OHCs and IHCs. Tbx2, Insm1, and Ikzf2 are critical for hair cell development, cell fate determination, and differentiation ^36–39^. The expression of these TFs needs to be tightly regulated to ensure their spatiotemporally expressing patterns to control programmed hair cell development and differentiation. Silencers ensure these powerful TFs are expressed only in designated cell types and not in others. However, after hair cells have differentiated and matured, these TFs are no longer critical for coordinated gene regulation. Instead, at later developmental or adulthood, they become cytoplasmic and may exert other functions beyond transcriptional roles. RFX family TFs have several natural splicing isoforms ^9,40^. The inconsistency of protein immunostaining and the RNA *in situ* data suggests that antibodies only recognized specific isoforms at different developmental stages. Rfx4 and Rfx8 have many natural isoforms, including those without a DBD domain^9^. The isoforms of Rfx3 and Rfx7 that show cytoplasmic localization may lack DBD domain or nuclear localization signals, adapting for a new function in the cytoplasm. Further investigation into other aspects of RFX TF functions beyond transcriptional regulation may be critical for discovering new mechanisms of hair cell homeostasis, maintenance, and immune response. This research will provide new insights into protecting hair cells from damages.

### The activation or repression roles of Rfx3-bound CREs depend on their sequence context and binding TFs

Group 1 RFX TF members (Rfx1/2/3) possess both transcription activation and repression domains (AD and DIM), enabling RFX TFs to act as dual-function regulators through these domains ^10^. However, the precise mechanism underlying this duality remains unclear. In this study, Rfx3 ChIP-seq, ATAC-seq, and scRNA-seq data collectively profile the regulatory features of Rfx3 target genes, revealing distinct regulatory features across different cell types. For example, Rfx3 may bind to the intronic CRE in *Vangl2* locus and repress its expression in hair cells or SGNs, as indicated by strong Rfx3 binding but weak or absence of ATAC-seq signals in this CRE of these cell types (Figure 6H). This CRE was previously validated to show enhancer activity in IHCs, surrounding supporting cells, and Greater epithelial ridge (GER) cells, but not in OHCs, surrounding supporting cells, or SGNs ^24^. The transgenic expression pattern of this Rfx3-bound CRE, inconsistent with the expression pattern of Rfx3 in the single-cell atlas or ATAC-seq peak feature, confirms the repressive roles of Rfx3 in transcription regulation. The results also revealed the expression activity of CRE in the *Vangl2* locus was cell type-specific, depending on the motif binding sites of TFs within the CRE. Single-cell atlas expression data show that TFs are differentially expressed in each cell type, such as Forkhead TFs in Figure 6. Thus, specific combinations of TFs in each cell type yield different regulatory outcomes, either activation or repression. The expression of Rfx3 and its binding to CRE may result in the repression of *Vangl2* gene specifically in hair cells and SGNs. In contrast, the absence of Rfx3 expression in supporting cells leaves the CRE of the *Vangl2* locus available for binding by other activating TFs, such as Six1 and Forkhead TFs (Foxp1) (Figure 6), which are reported to interact with RFX TFs and cooperatively activate target gene expressions ^24,30^. Therefore, activators and repressors compete to occupy the same CRE, with activity depending on the combination of binding TFs in different cell types. Our data revealed that Rfx3 primarily functioned as a transcriptional repressor and that its bound CREs can either be functionally activated or silenced, depending on their sequences context and interacting TFs.

Besides dimerization within family members via the DIM domain, the RFX TF family can also interact with other TFs to mediate combinatorial regulation of gene expression. RFX3 has been reported to interact directly with RFX1/2/4/6 and FOXJ1^30,41^. Our previous study demonstrated that Six1 cooperates with Rfx1/3 to potentially regulate the expression of *Pbx1* and *Dusp6* genes during inner ear development ^24^. Forkhead transcription factor Foxo3a interacts with RFX5, RFXB, and RFXAP, forming the MHC II enhanceosome and is essential for MHC II class gene transcriptions ^42^. In *Caenorhabditis elegans*, DAF-19, the only RFX TF identified, physically interacts with Forkhead TF FKH-8 to synergistically regulate ciliary gene expression ^43^. Our data identify the RFX motif in the Foxp1-bound CREs ^26^ and Forkhead motifs enriched in the Rfx3-bound CREs, suggesting interaction and cooperation between RFX and Forkhead TFs (Figure 6). Together, the dimerization and cooperation of RFX TF family members with other TFs are critical for orchestrating gene expression regulation, ultimately determining repression or activation.

### The redundant functions of the RFX family

Our data show that *Rfx3* and *Rfx7* are the two major RFX genes dominantly expressed in adult auditory hair cells (Figure 1). Previous studies revealed the redundant functions and compensatory effects between group 1 members, Rfx1-3, in mouse inner ears ^13^. Either single conditional knockout of *Rfx1* or *Rfx3* gene resulted in normal hearing, whereas compound Rfx1/3 cKO mice exhibited significant hearing loss ^13^. This suggests that the regulatory function of Rfx3 may be compensated by Rfx1 or other RFX TF members specifically expressed in hair cells. However, as Rfx1 expression is limited in mature auditory hair cells (Figure 1), its compensation is expected to be minimal. Therefore, Rfx7 is a strong candidate for compensating Rfx3. Rfx7 was reported to play roles in neural tube closure by regulating ciliogenesis in *Xenopus laevis* ^22^. It was also reported that Rfx7 plays critical roles in tumorigenesis, neurogenesis and immunity system ^19^. Many reported Rfx7 target genes identified in U2OS cell lines ^23^, such as *ABAT*, *ARL15*, *CABIN1*, *CDK4*, *JUNB*, *MXD4*, and *PNRC1*, were also found to be bound by Rfx3 in our mouse cochlea ChIP-seq data (Dataset S1). The common target genes of Rfx3 and Rfx7 further demonstrate their potential compensatory functions between each other. Previous study has shown the requirement of Rfx1/3 TFs for OHC survival in adult stages ^13^. Since Rfx1 is not the major RFX TFs expressed in OHCs, severe hearing deficiency is expected in Rfx3 and Rfx7 double cKO mice. Our project investigating Rfx7 function in mouse inner ear is currently undergoing.

The RFX TFs usually function as homodimers or heterodimers between family members. *In vitro* experiments revealed that Rfx1 can form dimers with Rfx2, Rfx3, and itself ^11^. The dimerization of RFX TFs is mediated by their DIM domain. Interestingly, motif analysis revealed that most of RFX motifs in the Rfx3 bound peaks in our study were also in “motif-dimer” form (two-half-site form) (Figure S3D). The DNA motif-dimer is expected to further enhance the dimerization of RFX TFs at the protein level. The presence of both DIM domain and motif-dimer suggests that the dimerization of RFX TFs is crucial for their functions and further reveals the cooperation and compensation between RFX TF members. However, Rfx7 differs significantly from Rfx3, belonging to different RFX TF family group (groups 1 and 3, respectively, Figure 1). Rfx3 and Rfx7 share very limited similarity, so their compensatory effect is expected to be limited due to their significant differences, despite being from the same family. The redundant and compensatory functions between Rfx3, Rfx7 and other RFX members remain unclear and require further investigation.

## Materials and Methods

### Animals

Wild-type C57BL/6 mice were bred in the laboratory. All animal experiments were conducted with approval from the Animal Research and Ethics Committee of the Chengdu Institute of Biology, Chinese Academy of Sciences (Approval No. CIBDWLL2018030).

### Chromatin immunoprecipitation sequencing (ChIP-Seq)

The Rfx3 ChIP-seq assay for inner ear tissue was conducted according to our previous protocol with minor modifications ^24,44^. Briefly, the cochlea epithelia were dissected from about 60 inner ears from wild-type E17.5 embryos. The dissected epithelia were cross-linked with 0.5% formaldehyde and 0.5% paraformaldehyde at room temperature for 2 hours, and then homogenized and lysed in cold lysis buffer (50 mM HEPES, pH 7.5, 140 mM NaCl, 1 mM EDTA, 10% glycerol, 0.5% NP-40, 0.25% Triton X-100, 1× protease inhibitors). Then, the nuclei were washed, pelleted at 2000 g at 4°C, and finally sheared into an optimal chromatin size in sonication buffer (10 mM Tris-Cl, pH 8.0, 2 mM EDTA, 0.1% SDS, and 1× protease inhibitors) using the Bioruptor Pico machine (Diagenode). The chromatin was precleared with Protein A Dynabeads (10002D, Invitrogen) and then incubated with 5 µl anti-RFX3 antibodies (HPA035689, Sigma) overnight at 4°C. The ChIP-seq with rabbit IgG antibody (PP64, Merck) was used as a control. The immunoprecipitated complexes were isolated by incubation with 20 µl new protein A Dynabead for 2 hours at 4°C, followed by a series of wash steps with low salt, high salt and LiCl wash buffer ^24^, and finally reverse crosslinking overnight at 65°C. The quality controls of ChIPed DNA were performed with Qubit 4.0 Fluoremeter (Q33238, Thermo Fisher Scientific) using dsDNA HS assay Kit (Q32854, Thermo Fisher Scientific). The sequencing libraries were prepared using the ThruPLEX DNA-seq Kit (R400674, Takara) and sequenced on the Illumina HiSeq 2500 system. The genomic input DNA was also used to prepare libraries and sequencing as controls for peaking calling.

### Peak calling, motif analysis, and gene otology enrichment analysis

The quality of the raw sequencing reads of ChIP-seq data was evaluated using FastQC (v0.11.9) (https://www.bioinformatics.babraham.ac.uk/projects/fastqc/). Raw reads were first trimmed using fastp with default parameters. The pair-end reads were aligned to mouse reference genome (mm10) using Bowtie2 with end-to-end parameter (v2.5.1) ^45^. Mapped reads were sorted using Samtools. Peak calling was conducted using MACS2 (v2.1.1) ^46^, and only the peaks with q value <0.05 were used for the following analyses. Peak annotation was performed using Chipseeker (v1.40.0) to retrieve the nearest genes around the peak and annotate genomic region of the peak (annotated by GENCODE vM23) ^47^. Motif enrichment analysis was conducted using the Homer package (v4.11) ^29^. Gene ontology analysis was performed using GREAT program ^48^.

### Immunofluorescence

The dissected cochlea was fixed in 4% paraformaldehyde (PFA) at 4°C for 2 hours. After fixation, the cochlea was washed, soaked in 30% sucrose overnight, embedded in OCT compound (4583, Sakura), sectioned at 12 μm. After air dry, the slides were washed and permeated in blocking buffer (containing 1×PBS, 5% BSA and 1% Triton X-100) overnight at 4°C. After blocking, the slides were incubated with primary antibody in buffer containing 1×PBS, 5% BSA, and 0.5% Triton X-100, at 4°C overnight. After washing with PBST for 5 min for 3 times, the slides were incubated with goat anti-Rabbit IgG Fluor 594 (111-585-003, Jackson) at 4°C overnight. Nuclei were counterstained with DAPI, and then the fluorescence signals were examined under fluorescent microscope (55i, Nikon). The combined protein immunofluorescence with RNA *in situ* detection used the similar steps.

### Constructs and transgenic enhancer reporter assay

The CREs bound by Rfx3 in *Triobp*, *Myo1h*, and *Insm1* gene loci were amplified from mouse genomic DNA as template using BamH1 and Xhol restriction enzymes. The primers are listed in Table S3. These CREs were cloned into the enhancer reporter vector with Hsp68 minimal promoter, pWhere-Hsp68-GFP ^24,44^. The constructs were linearized by PacI enzyme and purified for pronuclear injection. The injection was performed by Shanghai Model Organisms Center. The expression activity and specificity of these CREs were analyzed in transgene positive G0 embryos or P0 mice.

### RNA *in situ* hybridization

The gene fragments of *Rfx3*, *Rfx7*, *Triobp*, *Myo1h*, and *Ift81* were amplified from inner ear cDNA using the primers with the addition of T7 RNA polymerase site, as listed in Table S3. The antisense probes for each gene were synthesized and labeled using T7 RNA polymerase (P1440, Promega) and DIG labeled UTP (11277073910, Roche). The samples were fixed with 4% PFA for 5 min at room temperature (RT) and washed with PBS at RT for 3 min, then incubated in 0.2 M HCl for 10 minutes at RT. After being washed with PBS for 3 min, all slices were sequentially incubated with 0.1 M triethanolamine–HCl at RT for 10 min. After being washed in PBS at RT for 3 min, samples were dehydrated in a series of 75%, 85%, and twice in 100% ethanol at RT for 20min. Slices were then incubated in the hybridization buffer (50% formamide, 10 mM Tris-HCl, pH 8.0, 250 µg/ml yeast tRNA, 100 ug/ml Heparin and 1% SDS) containing probes at 60°C overnight. The slices were washed sequentially with 5×SSC at RT for 5 min, then with 2×SSC at 65°C for 30 min. All slices were treated with 0.3% H_2_O_2_ at RT for 1hour. After being washed twice with PBS at RT for 3 min, 200ul of blocking buffer (containing 1×PBS, 5% BSA and 1% Triton X-100) were dropped on the slide at RT for 1 hour, Finally, slices were incubated in blocking buffer with rabbit anti-DIG-POD (1:1,000; Roche) at 4°C overnight. Then the signal was detected by the TSA^®^ Plus Cyanine 3.5 (Cy3.5) detection kit (NEL763001KT, Akoya). Nuclei were counterstained with DAPI, before visualization on the fluorescent microscope system (55i, Nikon).

### Antibodies and Western blot

The rabbit anti-RFX antibody from Sigma was raised against 270-405 aa of human RFX3 protein (HPA035689, Sigma) and another anti-RFX3 antibody against 1-413 aa of human RFX3 were both used (14784-1-AP, Proteintech). The RFX7 antibody used in this study specifically recognized the 1313-1363 aa region of human RFX7 protein (A303-062A). Other antibodies used for Western blot or immunostaining were listed as followed: rabbit anti-Actin (AC026, Abclonal), mouse anti-Foxp1 (sc-398811, Santa cruz), HRP-conjugated goat anti-Rabbit IgG (H+L) (AS014, Abclonal), and goat anti-Mouse IgG Fluor594 (ab150116, Abcam).

For western blot, cochlear tissue was lysed in ice-cold lysis buffer (150 mm NaCl, 50 mm Tris-HCl, pH 8.0, 1 mm EDTA, 1% NP-40, 0.1% SDS) containing 1× protease inhibitor mixture (Roche), and precipitation was removed by centrifugation. Protein concentration was determined by BCA Protein Assay (BBCAPCK500, Bioswamp) and proteins were denatured at 100°C for 5 min in 1× Loading buffer (10% glycerol, 50 mM Tris-HCl (pH 6.8), 2% β-Mercaptoethanol, 0.02% Bromophenol blue, 2% SDS). Denatured proteins were separated by SDS-PAGE. The membrane was incubated with specific antibodies. After fully washed and then incubated with horseradish peroxidase (HRP)-conjugated secondary antibodies, the signals were developed using ECL reagent (WBKLS0500, Millipore).

### Data availability

The ChIP-seq and RNA-seq data in this study have been deposited in the National Genomics Data Center under the BioProject number (CAR017952). The ChIP-seq data of mouse inner ear from our previous publication (GSE108130 and GSE119545) ^24,44^, the public mouse sorted hair cell and SGNs ATAC-seq data (GSE181311) ^25^, and the Foxp1 ChIP-seq data (GSE101632) ^26^ were from public Gene Expression Omnibus database. The public mouse E16 and P7 stage inner ear single-cell RNA-seq data were accessed with the number GSE137299 ^49^. The mouse adult stage inner ear single-cell RNA-seq data have been deposited in the National Genomics Data Center under the number (CRA015540) ^27^.

## Acknowledgments

We thank Qian Liu from the Animal Experimental Center, College of Life Sciences, Wuhan University, for supporting in animal experiments. The work was supported by the National Natural Science Foundation of China (Nos. 82300142, 82071065, 82271187, 32370882, 82101233 and 32370536), Natural Science Foundation of Hunan Province (Grant No. 2022JJ30506 and 2022JJ30969), Startup Project from University of South China, and the University of South China Clinical Research 4310 Program, and the Central Government Guides Local Science and Technology Development Funds in Sichuan Provincial (2022ZYD0131).

## Author’s contribution

J. Li, Y. Feng, L. Ma and X. Guo designed the work; P. Zhang, Y. Wang, X. Guo, X. Zeng, Z. Feng, J. Liu, M. Yan, J.H. Li, and Y. Gao acquired the data; P. Zhang, Y. Wang, L. Ma, J. Ling and H. Wu, and Q. Jing analyzed the data; P. Zhang, Y. Wang, and J. Dong made the figures; J. Li, Y. Feng, L. Ma and G. Xiang wrote the paper.

