## Supplemental files for "Rfx3 controls outer hair cell differentiation, maintenance, and hair bundle formation by regulating the expression of *Insm1, Ikzf2*, and *Triobp* genes"

**Table S1: Ciliary genes bound by Rfx3**

| SYMBOL | Ciliary localization | distanceToTSS | Enrichment-fold | Disease or developmental defect |
|---|---|---|---|---|
| Kif3a | Cilia | -48 | 9.0 | Kidney cysts, retinal dystrophy, situs inversus (M) |
| Kif3b | Cilia | -31/448 | 122.4/2.5 | Situs inversus (M) |
| Ift88 | Cilia, basal bodies | 2 | 43.9 | Renal, hepatic and pancreatic cysts, hydrocephalus, polydactyly, situs inversus (M) |
| Ift172 | Cilia | -32 | 57.8 | Neural tube defects, polydactyly, situs inversus (M) |
| Ift122 | Cilia | 137/-2164 | 140.7/20.3 | Neural tube defects, polydactyly, situs inversus (M) |
| Ift140 | Cilia | -32/451 | 6.4/32.6 | Neural tube defects, polydactyly, situs inversus (M) |
| Ift22 | Cilia | -14 | 76.6 | Neural tube defects, polydactyly, situs inversus (M) |
| Ift27 | Cilia | 50 | 17.0 | Neural tube defects, polydactyly, situs inversus (M) |
| Ift43 | Cilia | 1 | 78.4 | Neural tube defects, polydactyly, situs inversus (M) |
| Ift74 | Cilia | 2 | 97.8 | Neural tube defects, polydactyly, situs inversus (M) |
| Ift80 | Cilia | 5 | 13.3 | Neural tube defects, polydactyly, situs inversus (M) |
| Ift81 | Cilia | -1 | 97.8 | Neural tube defects, polydactyly, situs inversus (M) |
| Ift52 | Cilia | -345 | 168.9 | Neural tube defects, polydactyly, situs inversus (M) |
| Ift57 | Cilia | -3 | 138.4 | Hydrocephalus, kidney cysts, situs inversus (Z) |
| Dnah5 | Cilia | -48838 | 80.6 | Sinusitus, bronchiectasis, infertility, hydrocephalus, situs inversus |
| Dnai1 | Cilia | -7 | 10.7 | Sinusitus, bronchiectasis, infertility, hydrocephalus, situs inversus |
| Dnah11 | Cilia | 94 | 106.7 | Situs inversus |
| Rfx3 | Nucleus | -5917/30 | 7.8/113.4 | Situs inversus (M) |
| Foxj1 | Nucleus | 1112/-37 | 3.8/229.3 | Situs inversus (M) |
| Pkd1 | Cilia, basal bodies | 563 | 55.7 | Kidney, liver and pancreatic cysts |
| Nphp1 | Cilia, basal bodies, centrosomes | 15 | 72.1 | Kidney cysts, liver fibrosis, retinal dysplasia |
| Nek1 | Cilia, basal bodies, centrosomes | 243 | 5.0 | Kidney cysts, male infertility; mouse model of progressive PKD |
| Smo | Cilia, cytoplasm, photoreceptor connecting cilium | 280/12713 | 3.8/8.0 | Neural tube defects, polydactyly (M) |
| Bbs1 | Basal bodies, centrosomes | -1 | 179.9 | Kidney cysts, obesity, anosmia, retinal dystrophy, male infertility, situs inversus, diabetes |
| Alms1 | Cilia, centrosomes | -1323/-45 | 3.8/31.2 | Retinal degeneration, obesity, diabetes |
| Mks1 | Cilia | -22 | 229.8 | Kidney and liver cysts, CNS malformations, polydactyly, hydrocephalus |

**Table S2: Hair bundle related genes bound by Rfx3**

| Gene Symbol | Protein | Peak position (bp-TSS) | Enrichment-fold | Human deafness |
|---|---|---|---|---|
| Diaph1 | Formin, GTPase-binding | 10627/1268/1730 | 56.7/63.3/15.1 | DFNA1 |
| Myo15a | MyTH4 ,FERM | -6021/-5463 | 7.0/2.9 | DFNB3\|MYO15 |
| Myo3a | Protein kinase | -308993 | 61.3 | DFNB30 |
| Myo6 | Myosin head, motor | -9572 | 2.5 | DFNA22\|DFNB37 |
| Ptprq | PTP type protein phosphatase | 57379 | 5.6 | DFNA73\|DFNB84 \|DFNB84A |
| Ush1c | PDZ | 2290 | 3.8 | DFNB18\|DFNB18A |
| Whrn | PDZ | -174 | 44.3 | DFNB31 |
| Cdh23 | Cadherin-like | 3126 | 7.8 | DFNB12 \| USH1D |
| Actg1 | Actin family | 31271 | 3.8 | DFNA20\|DFNB26 |
| Triobp | TRIO and F-actin-binding | 3334 | 11.4 | DFNB28 |
| Myo7a | Myosin head, motor domain | 61 | 6.4 | DFNA11\|DFNB2\|USH1B |

**Table S3: List of primer uesd in this study**

| Experiment | primer name | 5' to 3' sequence |
|---|---|---|
| Enhancer cloning | Insm1-enhancer-E17.5-BamH I-forward | CGCGCGGATCCATTGGCCAAAGCTGTCTCGG |
| Enhancer cloning | Insm1-enhancer-E17.5-Xho I-reverse | CGCCGCTCGAGCTGGCAAGAGACAACAACGAAGG |
| Enhancer cloning | Myo1h-enhancer-E17.5-BamH I-forward | CGCGCGGATCCGTGCAACACACCTACTCTCCT |
| Enhancer cloning | Myo1h-enhancer-E17.5-Xho I-reverse | CGCCGCTCGAGGCTGCATCTGTAACAACAGG |
| Enhancer cloning | Triobp-enhancer-E17.5-BamH I-forward | CGCGCGGATCCGAGGCTGTGTTTGTGGGAGG |
| Enhancer cloning | Triobp-enhancer-E17.5-Xho I-reverse | CGCCGCTCGAGCGGTAAGATCTAGGTGGGCC |
| RNA in situ | Ift81-RIS-forward | TGTTGATATCAGAGAGGAGATGCC |
| RNA in situ | Ift81-RIS-reverse | TAATACGACTCACTATAGTTCAGAACCTCAGTGCCGTC |
| RNA in situ | Myo1h-RIS-forward | TGGGAAAGCCTTCCGTGTTT |
| RNA in situ | Myo1h-RIS-reverse | TAATACGACTCACTATAGGGCCTTGAGCGTGGATAACT |
| RNA in situ | Rfx3-RIS-forward | TAGTCACCGTAGTCCTGGCG |
| RNA in situ | Rfx3-RIS-reverse | TAATACGACTCACTATATGTGACGCCCATCTGAATCC |
| RNA in situ | Rfx7-RIS-forward | TTGGCTGAATTCGTCCAGCA |
| RNA in situ | Rfx7-RIS-reverse | TAATACGACTCACTATAGCCCCTCTCCCATTTCACCAG |
| RNA in situ | Triobp-RIS-forward | CCTAGTCCTCGGTGTGTCCA |
| RNA in situ | Triobp-RIS-reverse | TAATACGACTCACTATAGAGGGGTATTGCTGTCTTGGC |

#### Figure S1

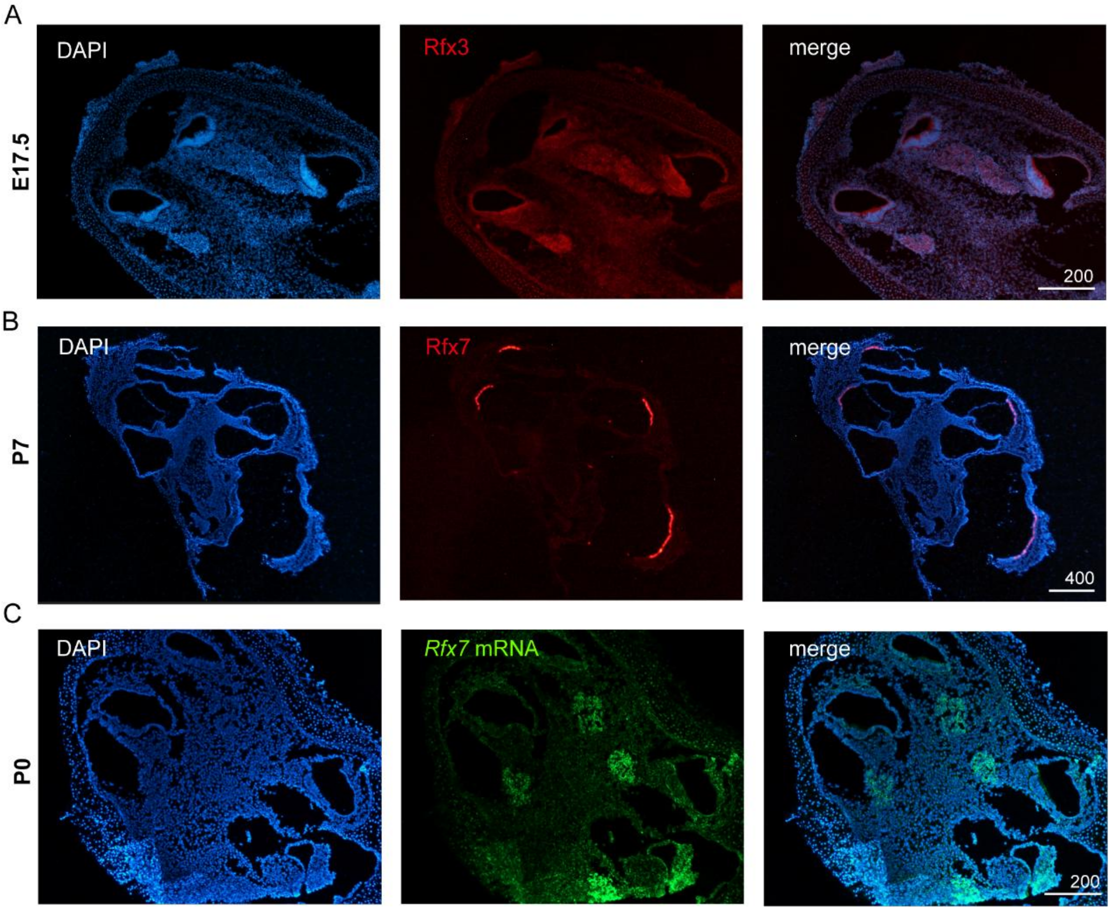

**Figure S1. Spatiotemporal and dynamic expression patterns of Rfx3 and Rfx7 in the mouse inner ear.** (A) Immunofluorescence detection of Rfx3 protein expression on cochlear sections at E17.5. Rfx3 immunostaining signals were labelled in red and nuclei were counterstained with DAPI in blue. (B) Immunofluorescence detection of Rfx7 protein expression on cochlear sections at P7. Rfx7 immunostaining signals were labelled in red and nuclei were counterstained with DAPI. (C) RNA *in situ* detection of *Rfx7* gene on cochlear sections at P0. Rfx7 mRNAs were labeled in green and the nuclei were counterstained with DAPI in blue. Scale bars are indicated in the figure (μm).

#### Figure S2

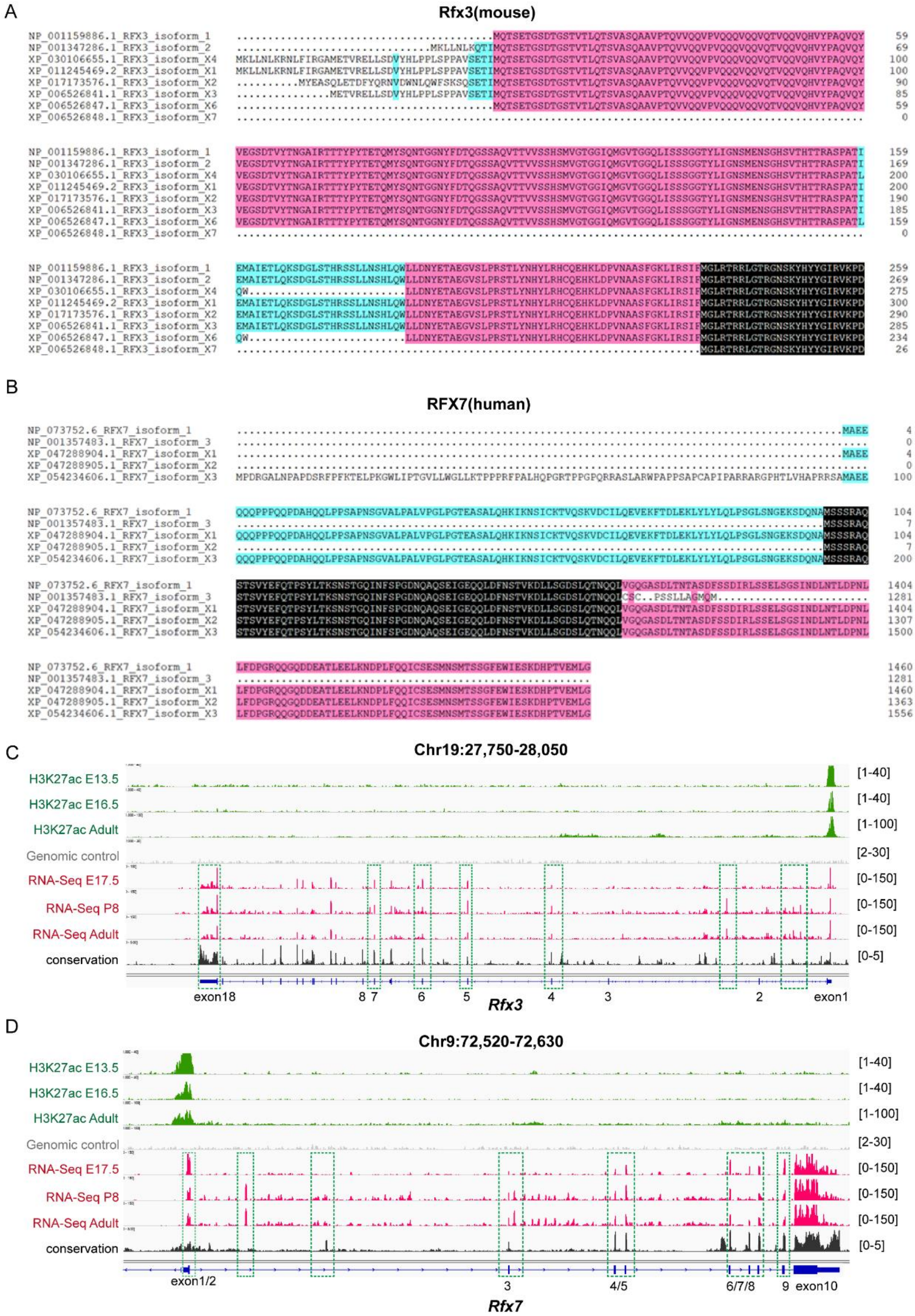

**Figure S2. Splicing and protein isoforms of Rfx3 and human RFX7 proteins**. Sequence alignments of Rfx3 isoforms (mouse) (A) and RFX7 isoforms (human) (B). Genome browser visualization of RNA-seq peaks at mouse Rfx3 (C) and Rfx7 (D) gene locus. The exons boxed by green dash displayed possible splicing variations.

### Figure S3

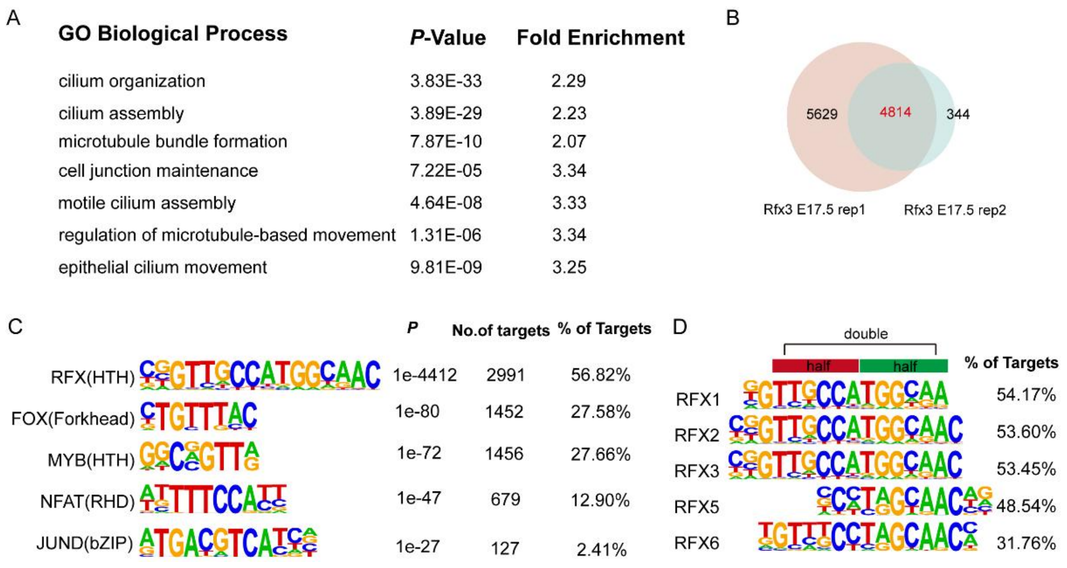

**Figure S3. ChIP-seq analysis of Rfx3 in mouse cochlea at E17.5 stage.** (A) The most enriched gene ontology (GO) terms of the Rfx3-bound genes in the replicate sample with fewer peaks in the replicate 2 sample. (B) The Venn diagram shows good overlap between two E17.5 ChIP-seq replicates. (C) The top 5 most enriched motifs in the Rfx3-bound CREs in the replicate 2 sample. (D) Sequence features of known X-box motifs from RFX family members. Either single or double half-site motif forms were identified according to the motif analysis of total Rfx3-bound CREs.

### Figure S4

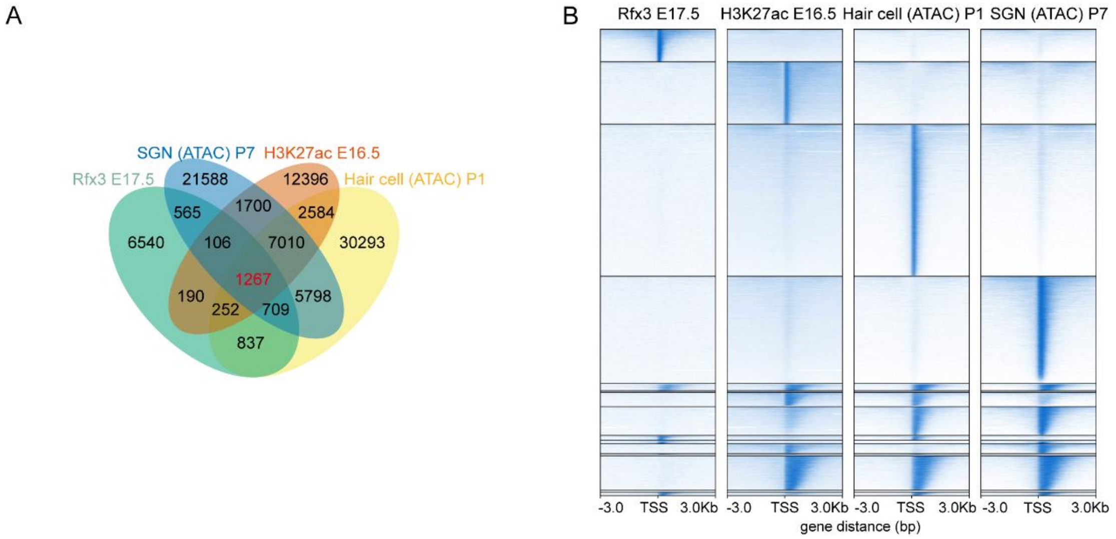

**Figure S4. Rfx3 may act either as a transcriptional activator or repressor.** (A) The Venn diagram shows the overlaps between Rfx3 binding site at E17.5, H3K27ac at E16.5, hair cell ATAC-seq at P1, and SGN ATAC-seq at P7. (B) Heatmaps showing the features of different clusters in Venn diagram in A, −3 kb/+ 3 kb window.

### Figure S5

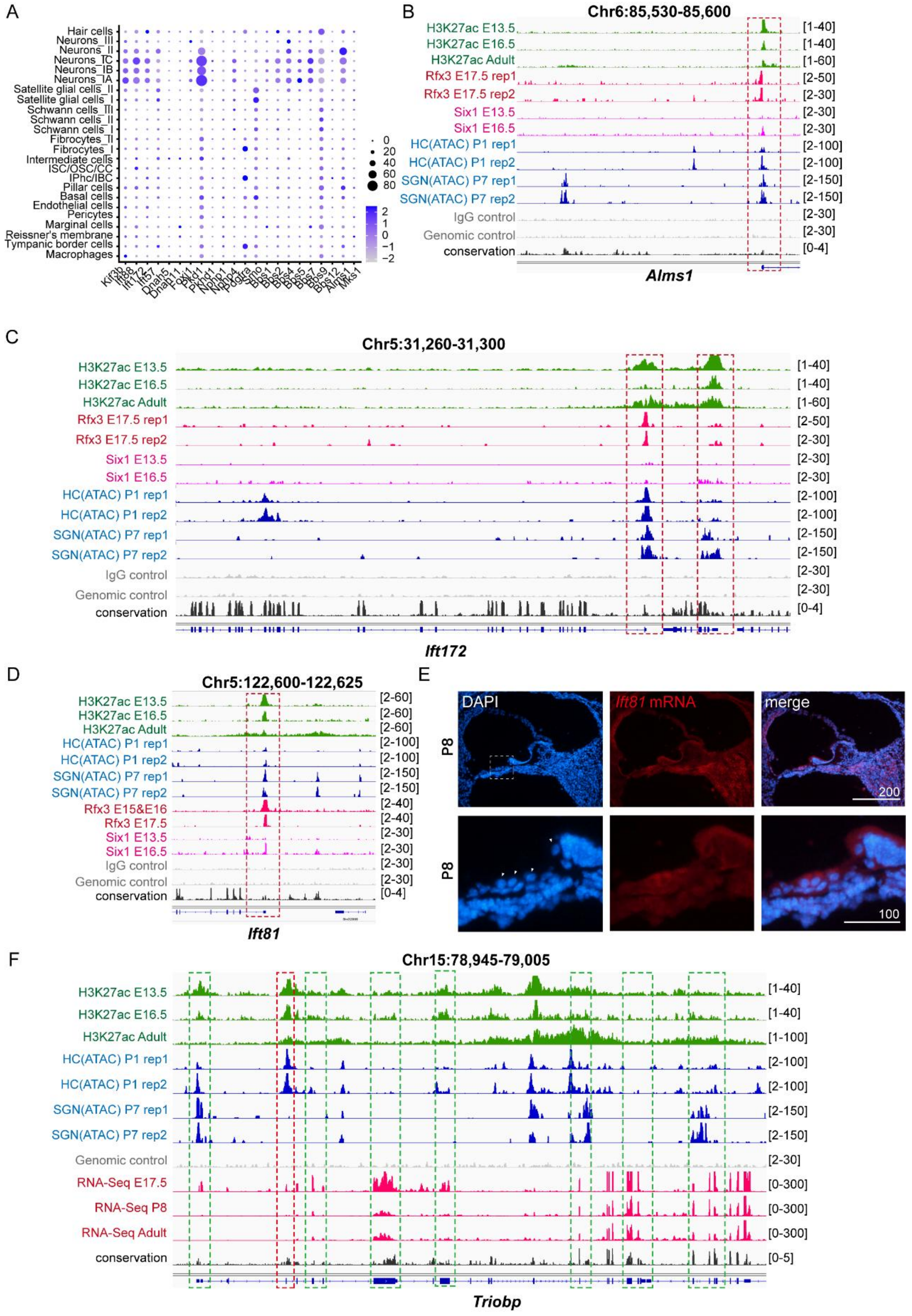

**Figure S5. Rfx3 regulates the spatiotemporal expression of ciliary and hair bundle related genes.** (A) Single-cell level profiling of ciliary gene expression in adult mouse cochlea. (B)

Genome browser visualization of Rfx3 peaks in *Alms1* locus. (C) Genome browser visualization of Rfx3 peaks in *Ift172* locus. (D) Genome browser visualization of Rfx3 peaks in *Ift81* locus. (E) RNA *in situ* detection of *Ift81* gene expression in organ of Corti at P8. The RNAs were labeled in green and nuclei were counterstained with DAPI (left panel, triangle indicates the position of hair cells), in blue, merge figure (right panel). (F) Genome browser visualization of RNA-seq peaks revealed the existence of splicing isoforms of *Triobp* gene. Scale bars are indicated in the figure (μm).

### Figure S6

A

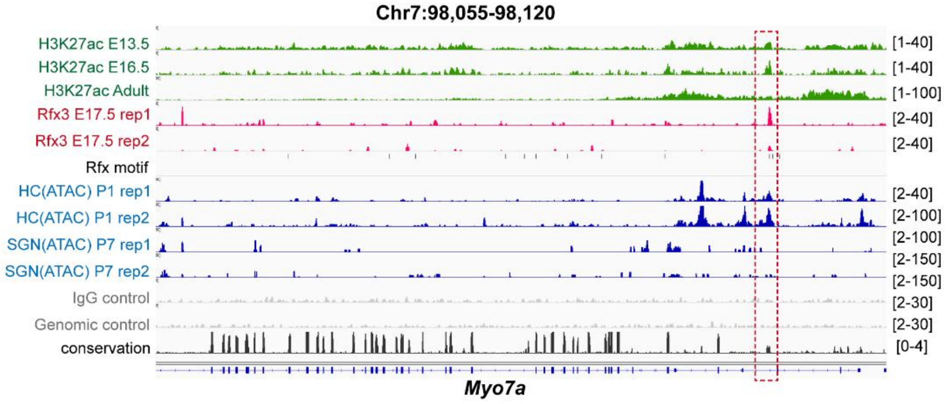

B

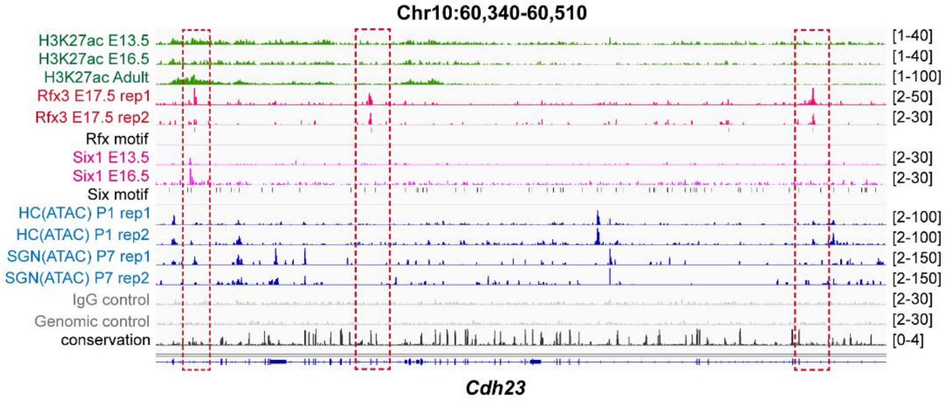

C

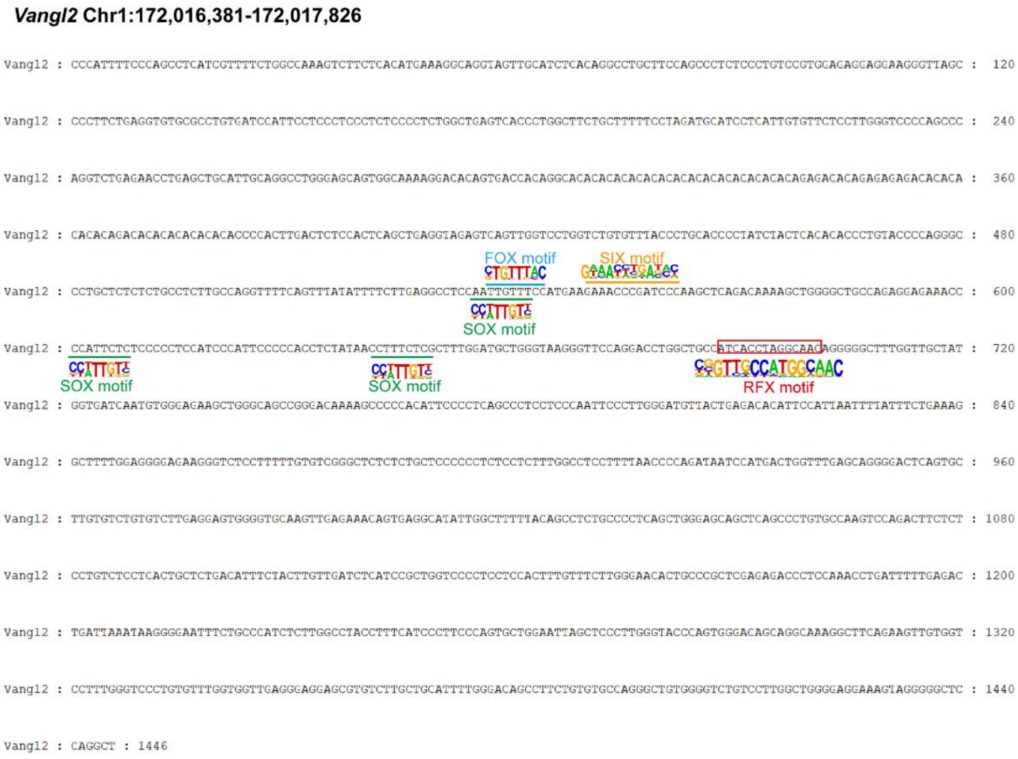

**Figure S6. Rfx3 regulates the expression of hair bundle genes *Myo7a* and *Cdh23*.** (A) Genome browser visualization of Rfx3 peaks in the intron of *Myo7a* locus. (B) Genome browser visualization of Rfx3 peaks in *Cdh23* locus. Three Rfx3-bound CREs were indicated by the red dashed boxes. (C) The *Vangl2* enhancer sequence contains several binding motifs of transcription factors critical for inner ear development.

#### Figure S7

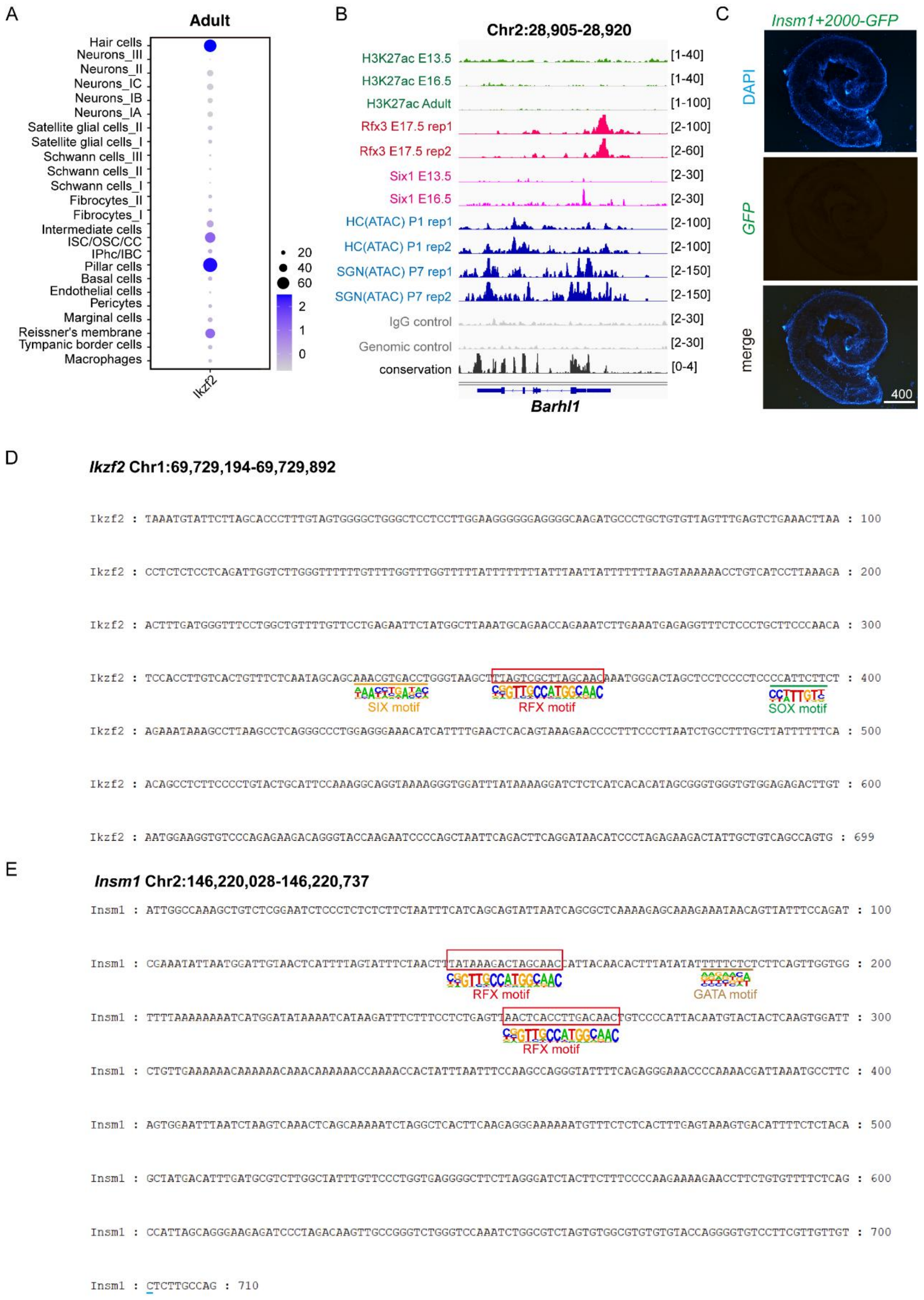

**Figure S7. Rfx3 regulates *Barhl1*, *Insm1*, and *Ikzf2* gene expression and plays critical roles in hair cell differentiation and maintenance.** (A) Expression of *Ikzf2* in the adult inner ear single cell atlas. (B) Genome browser visualization of Rfx3 peaks in *Barhl1* locus. (C) Transient (G0)

transgenic analysis of a 711-bp Rfx3-bound CRE in the *Insm1* intergenic region (the red-dashed-boxed-region in Figure 7C) at P0 stage. (D) Several critical TF motifs were marked in the Rfx3-bound CRE (Chr1:69729194-69729892) in *Ikzf2* locus, including RFX, SIX and SOX motifs. (E) The tested Rfx3-bound CRE (Figure7) in *Insm1* locus contains RFX and GATA motifs. Scale bars are indicated in the figure (μm).
